# High-Throughput Tracking of Freely Moving *Drosophila* Reveals Variations in Aggression and Courtship Behaviors

**DOI:** 10.1101/2025.07.10.663947

**Authors:** Saheli Sengupta, Ziying Chen, John Efromson, Aurélien Bègue, Mark Harfouche, Ayorinde S. Adegbesan, Amari E. Urquhart, Yick Bun Chan, Siyuan Yang, Sarah A. Henry, Caroline B. Palavicino-Maggio

## Abstract

Aggression is a nearly universal behavior used to secure food, territory, and mates across species, including the fruit fly *Drosophila melanogaster*. In fruit flies, both sexes display aggression through stereotypical motor patterns. This, along with their sophisticated genetic and molecular toolkit, makes *Drosophila melanogaster* an excellent model for studying aggression. While male- and female-specific aggressive motor programs have been qualitatively described, automated systems for quantifying these behaviors in freely moving flies remain limited in their ability to combine high-resolution analysis with high throughput. Here, we pair a high-resolution, high-throughput imaging system (the Kestrel) with DeepLabCut pose estimation to create a pipeline that tracks multiple freely moving fly pairs and quantifies social dynamics with high fidelity. We validated body-part tracking using published benchmarks. The platform reliably reproduced a known phenotype: heightened female aggression following thermogenetic activation of cholinergic pC1 neurons in female brain. It also detected increased unilateral wing extension, a courtship display inversely related to aggression, between two males upon activating a previously uncharacterized ∼40-neuron group in the male brain. Pose-based analysis revealed locomotive differences between experimental and control groups, and subtle, genotype-specific variations in head butts and UWEs. This workflow enables high-throughput screening and mechanistic dissection of social behaviors.

## Introduction

Social interaction, like aggression and courtship, is an evolutionarily conserved behavior important for securing essential resources, including food, territory, and mating partners. This behavior is highly sex-dependent: in *Drosophila melanogaster*, as in many species, males direct aggression toward other males while courting females as potential mating partners ^1–3^. This sex-specific interaction pattern reflects broader evolutionary strategies, in which dominant individuals – including those displaying high levels of aggression - gain access to limited resources, leading to increased survival and reproductive success. Given the strong influence of aggression and courtship on fitness, understanding these behaviors is essential for uncovering the evolutionary pressures and neural mechanisms that shape social interactions^1,4–6^.

Multiple vertebrate and invertebrate model organisms have provided valuable insights into the neurobiological and genetic basis of aggression and courtship. Among these, *Drosophila melanogaster* has emerged as an excellent model system because fruit flies display stereotypical, easily identifiable and quantifiable aggressive motor patterns. Their compact brain of approximately 200,000 neurons enables precise circuit-level analyses using an advanced genetic and molecular toolkit. In addition, many principles of circuit organization and genetic mechanisms are conserved between flies and mammals, including humans^7–11^. Despite morphological differences, the conservation of underlying genetic and neural mechanisms allows findings from fruit fly models to inform our understanding of aggression across species, including humans.

In *Drosophila melanogaster*, aggression is expressed through distinct motor programs: lunges in males and head butts in females. The male lunge involves a precise, multi-step action where one fly rises with its hind legs and snaps down on its opponent with its front legs. Female head butts involve striking an opponent with the head. Courtship, a reciprocal behavior to aggression, also involves stereotypical motor programs. A signature courtship behavior occurs when the courting male extends one wing at nearly 90° to his body and rapidly vibrates it, creating a courtship song that attracts mates. These simple but distinct behaviors provide a valuable framework for investigating the genetic and circuit mechanisms underlying social behaviors such as aggression and courtship^7,9,12–14^.

Many traditional imaging techniques for studying behavioral and neuronal dynamics require head-fixed or tethered animals. Two-photon microscopy provides high resolution imaging in deep tissue with specific neuron group targeting, but it also severely restricts natural movement and limits the range of observable behaviors. Quantification and definition of locomotive features of specific fly social interactions under free movement condition are still poorly explored^15–18^.

To address these limitations, closed-loop mechanical tracking systems have been developed to enable high-resolution imaging of freely moving subjects. While these systems have provided valuable insights into behavior and neuronal dynamics, they face several constraints. Most are limited to tracking a single organism at a time, making them unsuitable for studying interactions between multiple individuals. Additionally, these systems typically have a narrow field of view, limiting high-throughput applications^19–21^. Attempts to increase the field of view with large-diameter lenses introduce optical aberrations that require complex and expensive corrections^19,22,23^.

Fly behavior assays have been developed to increase throughput and consistency across experiments, while enabling accurate tracking of behavior^24,25^. These systems use video frame rates such as ≤30 Hz which work well for many behaviors but may present challenges when resolving rapid events like lunges or head butts, potentially leading to ambiguity in behavior annotation. Therefore, there is a need to build on these foundations by incorporating higher frame rates to capture fast behavioral events with greater temporal precision.

For post-experiment analysis, MATLAB-based tools such as FlyTracker^26^ and JAABA^27^ are commonly used for 2D position tracking and behavior classification. More recently, open-source deep learning tools like DeepLabCut^28–30^ and SLEAP^31^ have enabled more flexible pose estimation, greater pixel accuracy, and faster processing. To simplify the deep learning pose estimation implementation, some approaches were reported to constrain flies to a 2D space in camera view by limiting the vertical space within the behavioral arena. One such long-duration single-fly assay used vertical constraints to improve tracking accuracy, but this design could increase aggression in control group^24,32^. In some cases, siliconizing reagents have been applied to prevent flies from climbing on vertical surfaces perpendicular to camera view^33^.

To achieve high-throughput behavioral assay for investigating fly pair social interaction with their body parts tracked at high temporal and spatial resolution for a 30 min-assay time (used in this study), we customized a high throughput imaging system, the Kestrel, powered by the Multi Camera Array Microscope (MCAM) technology^19^, developed by Ramona Optics, Inc. This setup was paired with multi-view integration of three DeepLabCut pose estimation models from one view angle. The Kestrel is a cost-effective alternative to other large-area, high-resolution imaging systems, as it leverages inexpensive components commonly used in consumer electronics. The system provides high speed data acquisition in parallel from a grid of closely spaced cMOS image sensors (each with ∼10 million pixels) to reduce ambiguities due to slow acquisition cycles in behavioral recordings^19,34–36^. In our configuration, the Kestrel with a custom-designed chamber arena array uses 8 cameras to record high-resolution brightfield videos of multiple free moving fly pairs. To obtain position tracking from video recording, we implemented DeepLabCut, an open-source multi-subject pose estimation method based on convolutional neural networks (CNNs)^28–30^ and developed Python-based post-processing data analysis tools to integrate pose estimation output results from three DeepLabCut models based on fly body orientation relative to the camera angle. All videos were manually scored to assess levels of aggression and/or courtship between fly pairs.

We applied this system to investigate social behavior in genetically defined neural populations. Using the GAL4/UAS^37,38^ system, we thermogenetically activated specific neurons and examined changes in aggression and courtship. Using our setup, we found that activation of a small group of cholinergic pC1 neurons labeled by R26E01-GAL4 induced elevated female head butting, replicating previously reported results^9^. This confirmed that our imaging platform and behavioral setup reliably capture known behavioral outcomes. We next examined behavior in male–male assays by targeting neurons using R72A10-GAL4, a driver line that labels a previously uncharacterized group of ∼ 40 neurons in the central brain with few neurons in the ventral nerve cord.

Thermogenetic activation of these neurons induced a robust increase in unilateral wing extensions (UWEs) in temperature-activated R72A10 experimental males. UWE is a courtship display rarely observed in control male-male interactions^1,39^. This finding suggests that activation of R72A10-labeled neurons elicits enhanced male-male courtship without any effect on aggression. Furthermore, with the multiple pose estimation in individual fly with minimum no detection gaps at high resolution, our setup was able to measure basic features, like fly body segment size, and locomotive features about fly pair’s movement average across the whole recording session. From the manual scoring results, we measured locomotive features behind individual manually scored head butts and UWEs to investigate the difference in social behaviors behind elevated frequency of said behavior counts.

This integrated platform for high-resolution, high-throughput behavioral imaging offers a powerful new approach for quantifying aggression and courtship in fruit flies. By enabling precise tracking of body parts and behavioral outputs across multiple freely moving pairs, our system bridges a critical gap between manual scoring and scalable, automated analysis. Coupled with the genetic tractability of *Drosophila melanogaster*, this approach lays the groundwork for dissecting how specific neural populations shape distinct social behaviors.

Ultimately, such tools advance our ability to resolve fine-grained features of social behavior, link them to genetically defined circuits, and deepen our understanding of how aggression and courtship are regulated across species.

## Results

### Kestrel Fly Assay and Multi-View Integration of 3 DeepLabCut models from single camera view

To investigate social interactions such as aggression and courtship in *Drosophila melanogaster* at high resolution and across multiple parallel assays, we adapted the Kestrel system^19^ from Ramona Optics, Inc, Durham, NC (<u>Website</u>) (Fig. 1.a1). Behavioral assays were conducted using a customized base plate with 8 slots for cylindrical chambers (the fight arenas), which ensured consistent positioning for video recording across all experiments (Fig. 1.a2 and a3). This setup allowed simultaneous behavioral analysis of eight fly pairs. Each chamber has a removable glass ceiling that facilitates unobstructed imaging from the top. The cylinder’s internal dimensions (16.8 mm diameter, 7 mm height between floor and glass ceiling) allowed flies to move freely on both the floor and walls without spatial constraints that could artificially induce aggressive interactions. Within each chamber, a shallow recess (7 mm in diameter) holds 70 µL of standard fly food layered with 10 µL of yeast solution. A movable divider bisects each chamber, with small doors on either side that allow stress-free fly introduction using gravitaxis and phototaxis, eliminating the need for aspiration. This approach avoids extensive fly handling procedures such as aspiration or cold anesthesia, which have been reported to alter behavioral outputs by inducing stress^40^ (Fig. 1.a4).

**Figure 1.**
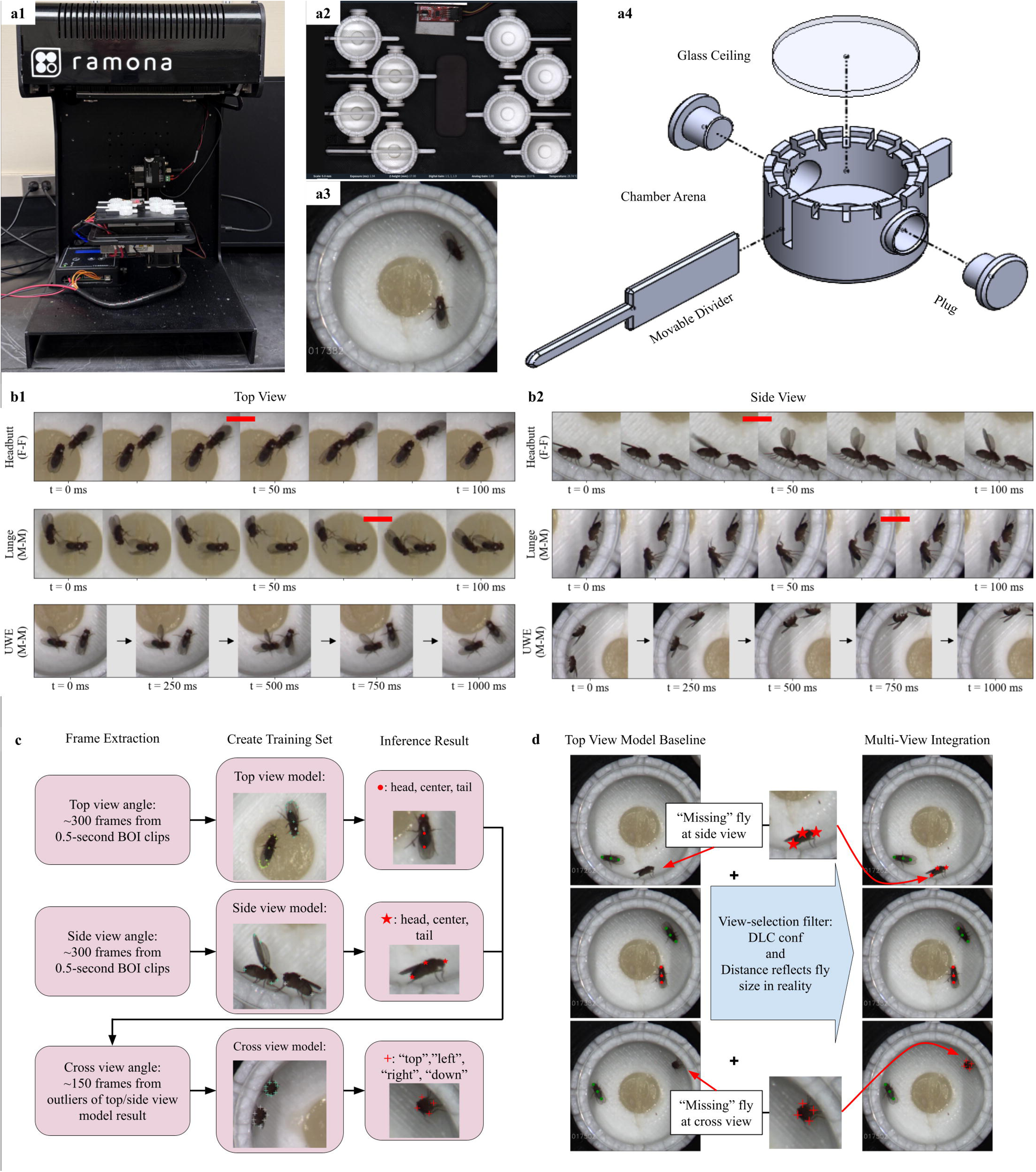
Multi-camera array setup and pose-estimation pipeline for recording parallel behavioral assay in *Drosophila melanogaster*. *a1*. Image of the MCAM-based imaging system used to record aggressive and courtship interactions across eight parallel chambers. *a2.* Top-down camera array view angle of 8-fight-arena setup: Left 4 chambers with dividers closed, right 4 chambers with dividers open. *a3.* Representative image illustrating how the MCAM Software automatically crops a single arena containing a fly pair after the experiment concludes. *a4.* Schematic of individual chamber, showing arena structure, glass ceiling, 2 plugs for fly transfer opening, and movable divider. *b1. and b2.* Manual annotations of behaviors of interest (BOI): head butts (HB), lunges, and unilateral wing extensions (UWE) and frame selections for pose estimation training. *b1.* Top view example: BOI when the fly pair is in dorsal-ventral body orientation. Top row and middle row show the frame sequence of a HB event in female-female (F -F) assay and a lunge event in male-male (M - M) assay at 60 frames per second (fps) respectively. Red bar indicates 2 consecutive frames with the largest travel distance in between during manual annotation. The bottom row shows the frame sequence of a UWE event in M-M assay at 4 fps. *b2.* Side view example: HB (top row), lunge (middle row) and UWE (bottom row) when the fly pair is in lateral body orientation. *c.* DeepLabCut (DLC) implementation: Frames are selected from 0.5-second clips of HB, lunge and UWE events in top view cases and side view cases. ∼300 frames in the top view were used for labeling and creating training dataset for the top view model, and the same is done for the side view model. ∼150 frames of fly pair in cross view body orientation, selected from outlier frames of top view and side view models’ inferencing results from 5 videos were labeled and used for training the cross-view model. Total number of body parts tracked in the three models: 12 in the top view model, 7 in the side view model, and 8 in the cross-view model. Body parts selected for locomotion analysis in this study: head, center, and tail in the top view model (circle) and the side view model (star), “top”, “left”, “right”, “down” in the side view model (cross). *d.* Multi-view integration of pose estimation models: The top-view model is the baseline, with red and green markers distinguishing the two flies. A view-selection filter was developed to fill the “missing” flies in cases where only one or even two flies were invisible in the top view, with side and cross view inferencing results.

We used Ramona’s MCAM software, the recording software for the Kestrel system, to capture fly-fly interactions at 30°C immediately after manually retracting the dividers. In our behavioral assays, we targeted different subsets of neurons in the fly brain using the GAL4/UAS binary expression system and expressed the thermosensitive cation channel TRPA1 in the GAL4-targeted neurons. It has been previously established that TRPA1 channel activates neurons at 30°C^38,41,42^. Each recording session captured all eight chambers simultaneously, and the first 30 minutes of each encounter between age-matched and sex-matched flies were analyzed. The MCAM software automatically crops out the physical space between arenas and generates eight videos at 576 × 576-pixel resolution at 60 fps for all chambers (Fig. 1.a3).

We focused on three key behaviors in *Drosophila melanogaster*: (1) male–male lunges, an aggressive motor program in which a male stands up on its hind legs and snaps down on the opponent with its front legs^7^ (Fig. 1.b1 and b2, middle); (2) male–male unilateral wing extension (UWE), a signature motor program of courtship behavior, in which one wing is extended at a right angle to the male body axis and vibrated^39,43–45^ (Fig. 1.b1 and b2, bottom); and (3) female–female head butts (HB), an aggressive behavior involving a forward thrust of the female head onto the opponent’s torso (Fig. 1.b1 and b2, top)^9,13,46^. Due to flies’ free movement in the custom chamber and single camera view angle of the Kestrel, a fly is usually shown in the top view when it is on the floor (Fig. 1.b1), in the side view when it is on the inner wall with body direction approximately horizontal to the floor (Fig. 1.b2), and in the cross view when it is on the inner wall with body direction approximately vertical to the floor when behavior cannot be properly observed to be manually scored. Hence, flies in two top or side view angles were considered when manually scoring for each behavior of interest (BOI).

In our experimental setup, observed lunges and head butts often span fewer than 3 frames at 60 fps, so during manual counting we define the event timestamp at the beginning of the two frames, indicated by the red bar in Fig. 1.b1 and b2, when the most displacement happens to the naked eye. UWEs happen on a slower timescale. In contrast to head butt and lunge events that are defined by rapid location change, UWE can last for hundreds of milliseconds and are defined by no change of specific wings’ orientation relative to the body. So, a UWE event timestamp is marked when the extended wing reaches its maximal angle, close to 90 degrees to the body direction. Each identified event is expanded into a 30-frame clip (i.e., 1/2 of a second) for downstream labeling and verification.

For pose estimation of multiple animals simultaneously, we implemented DeepLabCut (DLC)^28–30^. In order to minimize no-body-part-detection gaps for multi-animal pose estimation with almost no movement restriction, 3 distinct DLC models, with different labeled frames from manually scored event timestamps of BOI, were trained for 3 different viewing perspectives of the fruit fly in this experimental setup: top view, side view, and cross view (see details in “Methods”). As shown in Fig. 1.c. top and middle rows, key anatomical points - head, center, and tail-denoted by ● and Iii from top view and side view respectively, were used for locomotive feature analysis. The outlier frames from top view and side view models’ inference results were used to train the cross view model. Due to the unique circle shape when a fly shows its cross section in camera view, body points relative to the surface the fly is standing on, in Fig. 1.c. bottom row, were used to represent the fly coordinate in the cross view, with the point away from the surface as “top”, the part close to the surface as “down”, and the most left part of “top” and “down” as left, the most right point of the “top” and “down” as “right”. The coordinates of the four parts are averaged to calculate the approximate location, “cross”, as the fly body is in cross view orientation to the camera, often in its stationary posture.

As the baseline pose estimation for the fly pair, the top view model was chosen because dorsal-ventral body orientation is common for fly pairs in the custom fighting arena chamber. Pose estimation inference results from the side view model and cross view model are used to fill in “NaN” fly pose estimation from the top view model. A view-selection filter was developed to automate the process based on the DLC pose estimation confidence number (conf.) and distance between body parts reflecting average fly size. Fig. 1.d. showed a case where one fly in one frame is only detected by a side view model, then shows up in the top view model detection; later it is only detected by the cross view model pose estimation. The other fly is consistently being detected by the top view model inference. A Multi-View Coordinate Array (MVCA) is created to represent combined pose estimation, including head, center, tail, and “cross”, from all 3 models for the fly pair in 2-dimensional space across 30-minute duration. Figure 2. a. illustrated all the anatomical parts being tracked in the study by MVCA.

**Figure 2.**
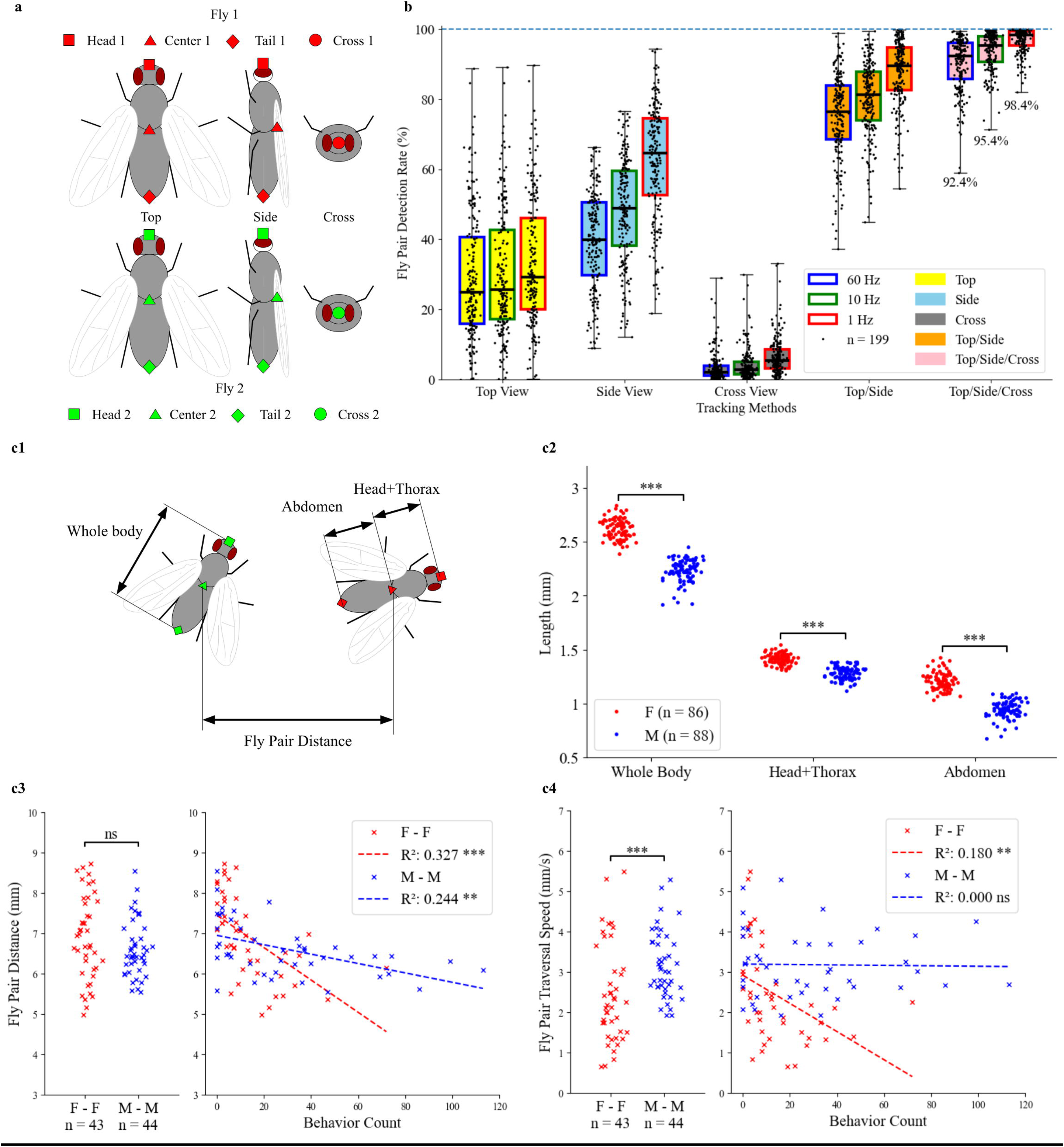
Fly pair detection accuracy and quantitative measures obtained from multi-view integration in MCAM fly-fly social behavioral assay. *a.* Body parts used in fly pair, fly 1 (green) and fly 2 (red), in different view angles for locomotion analysis. Head (square), center (triangle), and tail (diamond) in the top view and the side view; and cross (circle), an average location representation when a fly’s head or tail is towards the camera, in the cross view. *b.* Fly pair detection rate (%) comparison among different DLC pose estimation implementation in MCAM fly aggression experiment setup. Detection rates of 3 sampling rates for each method are presented, with 60 Hz as original recording frame rate, and 10 Hz and 1 Hz as downsampling results from original recording. The median fly pair detection rate of multi-view integration of top, side, and cross view is 92.4 % at 60Hz, 95.4 % at 10Hz, and 98.4% at 1Hz, across all the fly pairs, n = 199, in this study. Of all experiments, n = 86 were for studying fly pair with R26E01-GAL4 labeled neurons, n = 83 were for studying fly pair with R72A10-GAL4 labeled neurons, and n = 30 were for an independent group of TRPA1xW^1118^ fly pair. *c1.* Illustration for the definitions of fly whole body length, head+thorax length, abdomen length, and fly pair distance. In *c2.*, *c3.*, and *c4.*, TRPA1xW^1118^ fly pairs were used as the baseline to evaluate the tracking performance, including female-female (F - F) pairs, n = 43, and male-male (M - M) pairs, n = 44. As a result, for individual female flies (F), n = 86 and individual male flies (M), n = 88. *c2.* Length comparison of whole body, head+thorax, and abdomen length between individual female and male flies of TRPA1>W^1118^ line. For non-normal distributions via Shapiro-Wilk test, Mann-Whitney U test was performed for whole body and abdomen length comparison between female and male flies: ***, p < 0.0005 For normal distributions via Shapiro-Wilk test, independent t-test was performed for head+thorax length comparison between female and male flies: ***, p < 0.0005 *c3.* Left: fly pair distance (mm) comparison between female-female and male-male of TRPA1>W^1118^ line across 30 minutes. Independent t-test: ns, p ≥ 0.05. Right: scatter plot of fly pair distance vs. the manual count of behavior of interest, head butt for female and lunge for male. Linear regression: female-female, R² = 0.327, ***, *p* < 0.0005; male-male, R² = 0.244, **, *p* < 0.005. *c4.* Left: fly pair traversal speed (mm/s) comparison between female-female and male-male of TRPA1> W^1118^ line across 30 minutes. Mann-Whitney U test: ***, p < 0.0005. Right: scatter plot of fly pair traversal speed vs. the manual count of behavior of interest, head butt for female and lunge for male. Linear regression: female-female, R^2^ = 0.180, **, p < 0.005; male-male, R^2^ =0.000, ns, p ≥ 0.05.

### Evaluation of single DeepLabCut model and multi-view integration performance

To evaluate the DLC implementation and coverage of MVCA for freely moving fly pair social interaction during a 30-minute-long assay period, detection rate, fly physical size and some locomotive features were calculated from fly pair body parts inference results. First, we introduced the fly pair detection rate (see details in “Methods”), as a metric to quantify the share of duration of the experiment when multiple body parts were detected for both flies. The fly pair detection rates for all the fly pair videos in the study (n = 199) were shown in Fig. 2.b with different pose estimation implementations. For individual models, we can see that the top view model and side view model both had huge gaps of missing detection in their inference results, only covering less than 50% of the 30-minute recording session most of the time. Fly pairs both being in cross view was a rare occurrence. For combination models, by only combining top view and side view models, it dramatically improved the percentage of a recording session with viable fly pair detection. Additionally, with the cross view model, MVCA as a top/side/cross combination, has more than 90% fly pair detection rate for most of the fly pair videos. Besides at the original video recording frame rate, 60 Hz, which was essential for manual scoring of fast events like head butts and lunges, downsampling results of MVCA at 10 Hz and 1 Hz (see details in “Methods”) were used and evaluated in the study. Fig. 2.b showed that low sampling rate in general resulted in higher fly pair detection rate. Fly pair detection was used as a standard in all the locomotive feature calculations in the study. Furthermore, the event tracking rate was defined on top of the fly pair detection rate (see details in “Methods”) to measure the percentage of manually scored behaviors that had viable tracking results. For the three behaviors observed by experimenters in the study, head butt’s event tracking rate is 90%, UWE’s event tracking rate is 89%, and lunge’s event tracking rate is 93% (See Supplementary Data 1: *Event Tracking Rate*). This suggested the proximity and relatively high speed of fly pairs during fly interaction did not lead to selectively low tracking quality at timestamps around behaviors of interest.

To characterize the fly physical size and locomotive features from body part coordinates, female-female pairs (F–F, n = 43) and male-male pairs (M–M, n = 44) of the TRPA1xW^1118^ line in the study were used to benchmark the performance baseline. This genotype was used as a control line that carries the UAS-TRPA1 effector but lacks a GAL4 driver, thereby preventing thermogenetic neuronal activation at 30°C. This allows us to measure baseline physical and behavioral traits in the absence of neural activation. The limitation of a single model of top view or side view in coverage of the whole recording session does not affect its function to measure physical length of segments of a fly. A segment of a video, which meets certain threshold requirements, like continuous absence of non-detection, between-flies body parts distances, and relative location in the arena (see details in “Method”), was used to measure the whole body, head+thorax, and abdomen size of female and male flies as shown in Fig. 2.c1. The pose estimation was able to detect the general size difference between female and male. In Fig. 2.c2, estimated body segments of the females based on pose estimation from the top view model or side view model were larger than those of male counterparts, and result was consistent with a previous report about female and male body size comparison^47^.

For locomotive feature analysis, head butt counts in female-female pairs and lunge counts in male-male pairs were used as primary behavioral metrics because these two behaviors represent common aggressive interactions in control TRPA1xW^1118^ groups, in contrast to UWE events which are rare in male-male pairs and represent courtship, the reciprocal of aggression^1,7,13^. Fly pair distance is an average result across a 30-minute recording session using MVCA at the 60 Hz original recording rate, by using a single location to represent the coordinate of a single fly, then calculating distance between two flies using the single coordinate for each one (see details in methods). In the left panel of Fig. 2.c3, no significance difference in fly pair distance was found between female-female pairs and male-male pairs and in the right panel, both female-female pairs and male-male pairs showed negative correlation between fly pair distance and respective behavior count, head butt for females (*R*^2^ = 0.327, *p* < 0.0005) and lunges for males (*R*^2^ = 0.244, *p* < 0.005).

Fly pair traversal speed was a measure of speed that treats a fly pair as a whole to circumvent the fly identity switching in pose estimation (see details in “Methods”). MVCA at 10 Hz sampling rate was used because the purpose of the measurement was to assess the travelling distance that the fly pair covered in the arena, not localized fast movement during flies’ interactions. A lunge or head butt would not surpass the time window of 6 frames at 60 fps, so MVCA at 10 Hz was chosen. In the left panel of Fig. 2.c4, male-male pairs traversed faster as a whole than female-female pairs during the 30-minute recording session, consistent with the reported finding that male pairs had increased movement in confined space after maturing in isolation^48^. In the right panel, male-male pairs did not show correlation between their fly pair traversal speed and lunge count (*R*^2^ = 0.000, *p* ≥ 0.05) but female-female pairs showed negative correlation between their fly pair traversal speed and head butt count (*R*^2^ = 0.180, *p* < 0.005), which was a weaker correlation than those observed for fly pair distance vs behavior count in both female-female pairs and male-male pairs.

To further evaluate the setup’s potential to detect aggressive patterns (e.g., high count of head butts) and courtship patterns (e.g., enhanced UWEs) that a fly pair displayed, thermogenetic activation experiments of R26E01-GAL4 and R72A10-GAL4 driver lines via TrpA1 were conducted. Manual behavioral scoring—often considered the gold standard for behavioral analysis^24^ - and basic locomotion measurements were used to demonstrate the system’s capability to replicate previously reported results and make novel findings in fruit flies.

### Validation of R26E01-GAL4 female aggression and novel locomotive feature analysis using the Kestrel system

Previous research identified a small cluster (approximately 2–4 pairs) of *dsx+* cholinergic neurons situated near the pC1 neuronal cluster, targeted by the R26E01-GAL4 driver line. Thermogenetic activation of these neurons via TrpA1 (referred to here as R26E01>TRPA1) at 30 °C was previously shown to significantly enhance head butts, a type of female specific aggression^9^. In this study, we aimed to test whether our customized Kestrel system, combined with the newly designed behavioral arena, could be used to conduct a behavioral assay with fruit flies and successfully replicate the previously reported aggression phenotype resulting from thermogenetic activation of R26E01 neurons. To do so, we targeted the aforementioned neuronal cluster group using the R26E01-GAL4 driver line, performed thermogenetic activation at 30 °C as previously described^9^, and quantified the frequency of head butts in female-female dyadic encounters. In the 30-minute assay time, female pairs with activated R26E01 neurons indeed resulted in significantly elevated head butt frequency compared to female pairs in R26E01 control and TRPA1 control groups (Fig. 3.a. left panel; Supplementary video 1). This replicated the previously observed aggression phenotype and demonstrated that our Kestrel system and arena setup can reliably be used to study social behaviors such as aggression in fruit flies.

**Figure 3.**
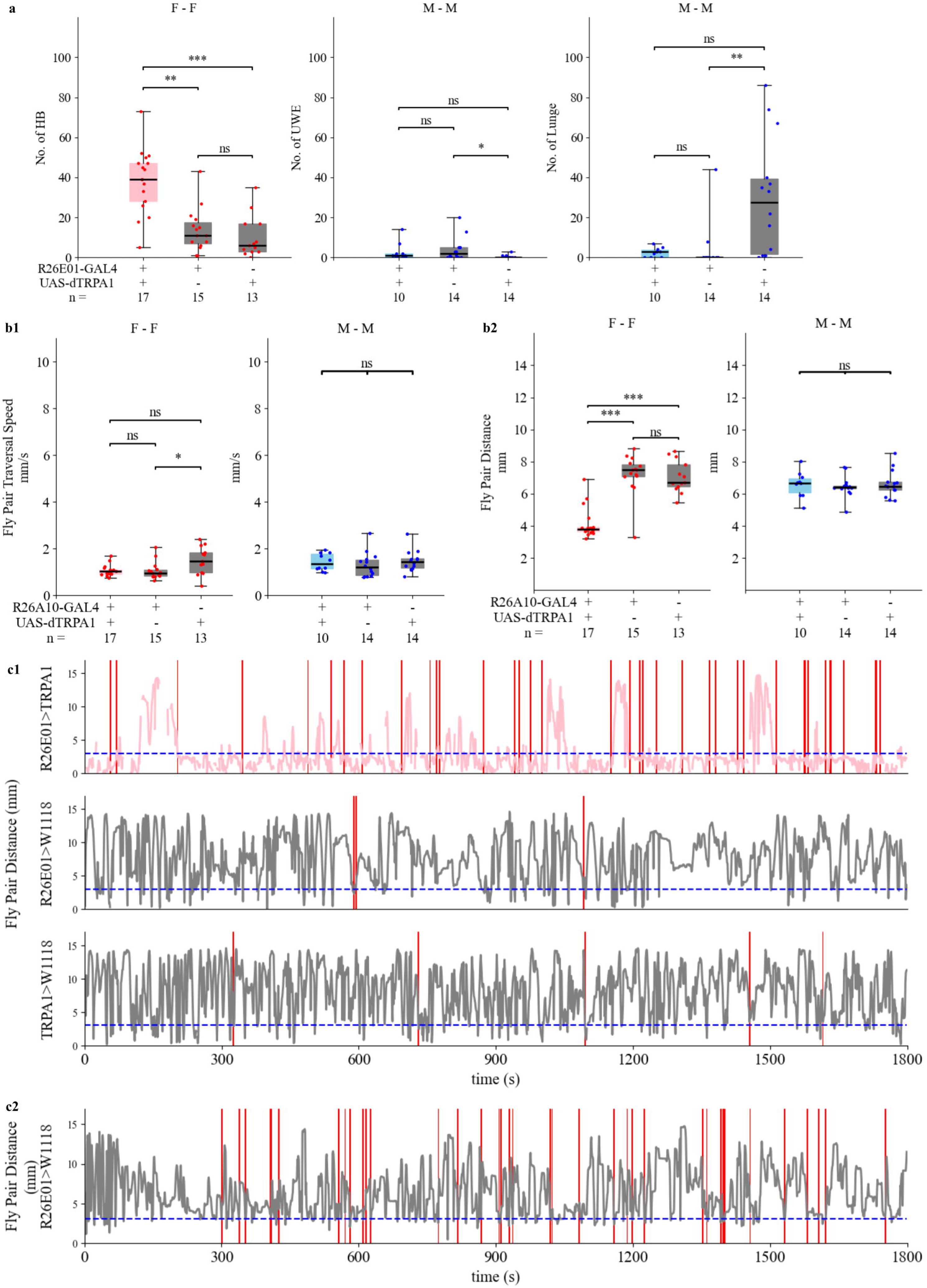
Thermogenetic activation experiment of R26E01-GAL4 labeled neurons: Manual score of head butt, lunge, and UWE, and locomotion from fly tracking. *a.* Manual score of behaviors of interest for 3 genotypes: R26E01>TRPAI (experimental), R26E01>W^1118^ (control), and TRPA1>W^1118^ (control). Left: head butt counts in female–female pairs (n = 13–17). Center: UWE counts in male-male pairs (n = 10–14). Right: lunge counts in male-male pairs (n = 10–14). Kruskal–Wallis test was performed for multiple comparisons within each behavioral category. Significance was observed for all behaviors of interest (*p* < 0.05, Kruskal–Wallis test). *b1. and b2.* Locomotion from pose estimation comparison among 3 genotypes: R26E01>TRPAI (experimental), R26E01>W^1118^ (control), and TRPA1>W^1118^ (control). *b1.* Fly pair traversal speed (mm/s) comparison among 3 genotypes. Left: female-female assay (*p* < 0.05, Kruskal– Wallis test). Right: male-male assay (ns, *p* ≥ 0.05, Kruskal–Wallis test). *b2.* Fly pair distance (mm) comparison among 3 genotypes. Left: female-female assay (*p* < 0.05, Kruskal–Wallis test). Right: male-male assay (ns, *p* ≥ 0.05, Kruskal–Wallis test). For all three-genotype comparisons in *a.*, *b1.*, and *b2.*, post-hoc pairwise comparisons were performed using Dunn’s test with Bonferroni-adjusted *p*-values: ***, *p* < 0.0005; **, *p* < 0.005; *, *p* < 0.05; ns, *p* ≥ 0.05. Pink box: experimental group of female-female. Blue box: experimental group of male-male. Gray box: control groups of the sex in comparison. Red dot: individual female-female pair. Blue dot: individual male-male pair. *c1.* Representative 30-minute assay of female-female interaction showing head butt events and corresponding changes in fly pair distance (mm) over time. Top: R26E01>TRPA1, Middle: R26E01>W^1118^, and Bottom: TRPA1>W^1118^. The pink and grey traces represent fly pair distance over time in experimental (c1 top) and control flies (c1 middle, and c1 bottom), respectively. *c2.* Representative 30-minute assay of an atypical female-female interaction in the R26E01> W^1118^ control group, showing elevated head butt events (red ticks) without sustained periods of short fly pair distance (gray trace, in mm) over time. In *c1.* and *c2.*, the x-axis represents time in seconds. Vertical red bars mark head butt events, with thicker red bars indicating bursts of high-frequency head butts within a short time window. The horizontal dashed blue line marks 2.5 mm, which approximates the whole-body length of a fly and serves as a threshold indicating proximity between flies.

In addition, we also measured basic locomotive features, fly pair traversal speed and fly pair distance during the 30-minute assay time. The increased head butt counts in the R26E01>TRPA1 group did not reflect greater traversal speed for the female pair as whole compared to control groups, shown in Fig. 3.b1 left panel, but resulted in significantly closer fly pair distance over 30 minutes in Fig. 3.b2 left panel.

Here, we also showed the time courses of head butts and fly pair distance for female pairs with tracking at 1 Hz sampling rate in Fig. 3.c1, with one example from each genotype. Besides the aggression represented by high head butt count in the top panel of Fig. 3.c1, the example female-female pair with temperature-activated R26E01 neurons was shown to have persistent periods of extremely close fly pair distance, shorter than 2.5mm, a common body length of a female fruit fly measured by this setup. We showed the time course of a female pair with a manual count of head butt events in the R26E01 control group in Fig. 3.c2. Despite the similarity of head butt frequency to the experimental R26E01>TRPA1, the fly pair distance trace of the R26E01 control group pair was similar to the other two control examples in the middle and right panels of Fig. 3.c1. Overall, these results demonstrate that the Kestrel and the newly designed behavioral chamber successfully replicate previously observed findings and provided fly pair distance, an additional locomotive measurement that indicates that the R26E01-activated females persistently stay in close distance

Next, we extended our analysis to evaluate male-specific behaviors using the same genotypes. Consistent with previously published observations, thermogenetic activation of R26E01 neurons did not significantly alter the frequency of lunges in male-male interactions in Fig. 3.a middle panel. Additionally, we analyzed the same male-male assay videos to score UWEs, which are commonly recognized as courtship “love songs” and are behaviors reciprocal to aggression. This feature was not analyzed in the previous study^9^, and our analysis provides additional insight into the behavioral repertoire of R26E01>TRPA1 males. Our analysis revealed no statistically significant difference in the frequency of UWEs between R26E01>TRPA1 males and control males. Similar to the controls, the frequency of UWEs in R26E01>TRPA1 male–male assays was low.

Additionally, male–male pairs of the three genotypes— R26E01>TRPA1, R26E01 control, and TRPA1 control**—** did not show a significant difference in fly pair traversal speed or fly pair distance over 30 minutes, as shown in the right panels of Fig. 3b1 and Fig. 3b2. Collectively, these results confirm that thermogenetic activation of R26E01 neurons selectively modulates aggression in females, without significantly impacting male aggression, courtship behaviors, or locomotor features. Furthermore, body part tracking measurements derived from multi-view DeepLabCut (DLC) integration were consistent with the manually observed behavior counts. Given the reliable reproduction of previously reported aggression findings, we then used the Kestrel system to conduct systematic behavioral studies in additional GAL4 fly lines.

### Investigation of social behavior and locomotive features in R72A10-GAL4 driver line using the Kestrel system

In a screen from a previous study^49^ for identifying neural substrates of aggression, we examined the R72A10- GAL4 driver line and found that thermogenetic activation of the R72A10-GAL4 labeled neurons at 30**°**C did not significantly alter aggression levels in male–male assays. Repeating this experiment with the Kestrel system yielded consistent results, with no change in aggression in the temperature-activated R72A10>TRPA1 experimental males at 30**°**C relative to controls. However, we found that during the 30-minute assay time between two males, the experimental R72A10>TRPA1 males exhibited a substantially increased frequency of UWEs, but not lunges, compared to controls (Fig. 4a, middle and right panels; Supplementary video 2). In wild-type males, aggressive behaviors dominate such interactions, and courtship displays such as UWEs are typically absent or rare^1,39,49,50^. The robust induction of UWEs upon thermogenetic activation of R72A10-GAL4 neurons suggests that these neurons elicit enhanced courtship in male-male encounters without any effect on aggression. Previous studies have shown that silencing a cluster of GABAergic neurons, popularly known as mAL and situated medially above the antennal lobes, enhances UWEs in male-male encounters.^49^ These findings raise the possibility that R72A10-GAL4 targeted neurons interact with mAL neurons in a circuit and functionally inhibit them, resulting in increased UWEs in male-male encounters. Future experiments will be necessary to test this hypothesis.

**Figure 4.**
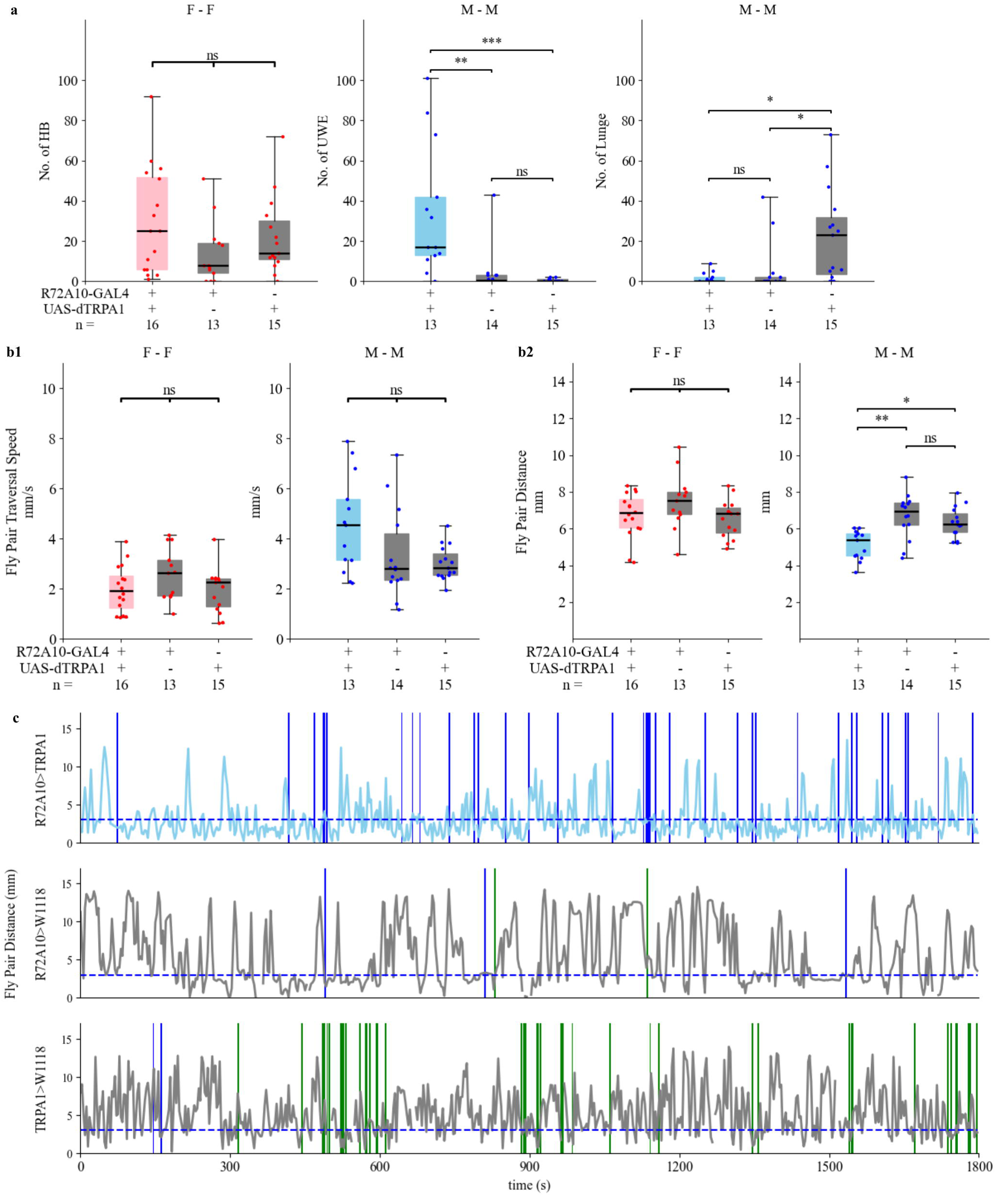
Thermogenetic activation experiment of R72A10-GAL4 labeled neurons: Manual score of head butt, lunge, and UWE, and locomotion from fly tracking. *a.* Manual scoring of behaviors of interest for 3 genotypes: R72A10>TRPA1 (experimental), R72A10> W^1118^ (control), and TRPA1> W^1118^ (control). Left: head butt counts in female–female pairs (n = 13–16). Center: UWE counts in male-male pairs (n = 13–15). Right: lunge counts in male-male pairs (n = 13–15). Kruskal–Wallis test was performed for multiple comparisons within each behavioral category. No significant difference was found in head butt counts in female-female pairs across 3 genotypes (*ns*, *p* ≥ 0.05), while both UWE and lunge counts in male-male pairs showed significant differences across 3 genotypes (p< 0.05, Kruskal–Wallis test). *b1. and b2.* Locomotion from pose estimation comparison among 3 genotypes: R72A10>TRPAI (experimental), R72A10>W^1118^ (control), and TRPA1>W^1118^ (control). *b1.* Fly pair traversal speed (mm/s) comparison among 3 genotypes. Left: female-female (ns, *p* ≥ 0.05, one-way ANOVA test). Right: male-male (ns, *p* ≥ 0.05, Kruskal–Wallis test). *b2.* Fly pair distance (mm) comparison among 3 genotypes. Left: female– female (ns, *p* ≥ 0.05, one-way ANOVA test). Right: male–male (*p* < 0.05, one-way ANOVA test). In *a.*, *b1.*, and *b2.* for 3-genotype comparisons that passed one-way ANOVA test, Tukey’s test was performed with adjusted *p*-values: ***, *p* < 0.0005; **, *p* < 0.005; *, *p* < 0.05; ns, *p* ≥ 0.05. For 3-genotype comparisons that passed Kruskal-Wallis test, post-hoc pairwise comparisons were performed using Dunn’s test with Bonferroni-adjusted *p*-values: **, *p* < 0.005; *, *p* < 0.05; ns, *p* ≥ 0.05. Pink box: experimental group of female-female. Blue box: experimental group of male-male. Gray box: control groups of the sex in comparison. Red dot: individual female-female pair. Blue dot: individual male-male pair. *c.* Representative 30-minute assay of male-male interaction showing UWE and lunge events and corresponding changes in fly pair distance (mm) over time. The x-axis represents time in seconds. Top: R72A10>TRPA1, Middle: R72A10> W^1118^, and Bottom: TRPA1> W^1118^. Vertical green bars mark lunge events, and vertical blue bars mark UWE events, with thicker green or blue bars indicating bursts of high-frequency lunge or UWE within a short time window. The blue and grey traces represent fly pair distance over time in experimental (c top) and control groups (c middle, and c bottom), respectively. The horizontal dashed blue line marks 2.5 mm, which approximates the average whole-body length of a fly and serves as a threshold indicating proximity between flies.

Next, we measured basic locomotor features. The high frequency of UWEs observed in the R72A10>TRPA1 experimental group did not reflect a significant change in traversal speed of the R72A10>TRPA1 male pair compared to controls (Fig. 4b1, right panel). However, it resulted in a significantly closer fly pair distance over the 30-minute assay time (Fig. 4b2, right panel). We also performed time-course analyses of lunges, UWEs, and fly pair distance for experimental and control male–male assays using tracking at a 1 Hz sampling rate. In the representative assay for the R72A10>TRPA1 experimental flies, in addition to high UWE counts (top panel, Fig. 4c), the males exhibited persistent periods of very close fly pair distance.

Using the Kestrel system, we uncovered a previously uncharacterized neuronal population targeted by R72A10-GAL4 driver line whose activation selectively promotes male courtship behavior, laying the foundation for dissecting the circuit-level mechanisms that govern social decision-making.

We also investigated aggression in R72A10>TRPA1females at 30**°**C. Manual scoring of head butts in female-female dyadic encounters (Fig. 4.a. left panel) revealed no statistically significant difference between experimental and control groups. Female pairs of the three genotypes—R72A10>TRPA1, R72A10 control, and TRPA1 control—also did not show a significant difference in fly pair traversal speed or fly pair distance over 30 minutes (left panels of Fig. 4b1 and Fig. 4b2).Together, these findings highlight the sex-specific behavioral effects elicited by activation of R72A10 neurons and underscore the utility and reliability of the Kestrel in discovering novel neuronal functions in behavior.

### Aggression and pursuit indices as strong indicators of social behaviors in thermogenically activated R26E01>TRPA1 and R72A10>TRPA1 flies

Basic locomotive features, like fly pair traversal speed and fly pair distance, were calculated using a single coordinate representing the location of a single fly. Fly pair traversal speed over 30 minutes showed little to no correlation to head butt counts of female-female interactions (Fig. 2.c4, Fig. 3.b1 and Fig. 4.b1). Fly pair distance over 30 minutes was a strong general indicator of increased behaviors, either head butts between females or UWEs and lunges between males (Fig. 2.c3, Fig. 3.b2, and Fig. 4.b2). In Fig. 5.b, the fly pair distance is presented here as an indicator to compare all 6 genotype groups that we recorded in R26E01-GAL4 targeted- and R72A10-GAL4-targeted-neuron activation experiments. We found that the fly pair distance over 30-minute encounter was a very strong indicator of elevated head butt frequency in R26E01>TRPA1 activated female pair compared to the other 5 genotypes (Fig. 5.b left panel). However, enhanced courtship in R72A10>TRPA1 activated male pair did not result in a significantly close fly pair distance when compared to the control groups.

**Figure 5.**
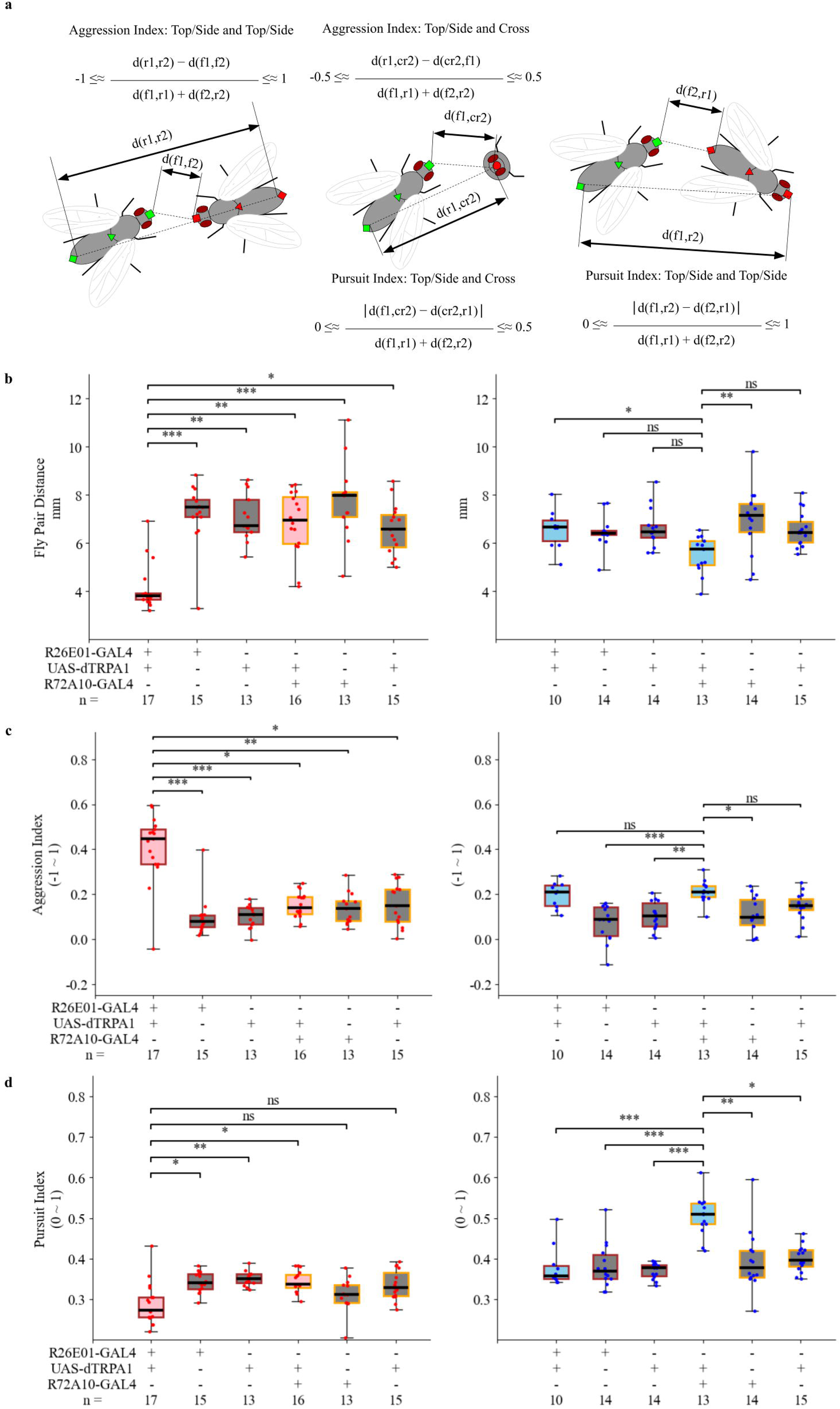
Definition of aggression and pursuit index and comparison of fly pair distance, aggression index and pursuit index during 30-min experiment time across different genotypes. *a.* Illustration of aggression index and pursuit index definitions based on view angles of the fly pair. Each index is derived from distance measurements between paired body landmarks of two flies, then normalized by body part length. f1, f2: front parts and r1, r2: rear parts of selected two body parts from the two flies, cr2: cross of fly 2 in the cross view representation, d: Euclidean distance of two tracked body parts. Left: aggression index when both flies are visible in top or side view. Its value ranges from –1 to 1 approximately. The higher value indicates that the two flies tend to be in head-to-head position during the experiment. Right: pursuit index when both flies are in top or side view. Its value ranges from 0 to 1 approximately. The higher value indicates that two flies tend to be in chasing or following body position during the experiment. Center: aggression index and pursuit index when one fly is in top or side view and the other is in cross view. Due to limitation of cross view model detection, aggression index value range in this situation is approximately from –0.5 to 0.5, and pursuit index value range is approximately from 0 to 0.5. *b.* Fly pair distance (mm) across different genotypes. Left: fly pair distances in female-female assays across different genotypes. R26E01>TRPA1 female pairs showed significantly shorter fly pair distance compared to the other five female groups. Right: fly pair distances in male–male. *c.* Aggression index across different genotypes. Left: aggression index in female–female assays. R26E01>TRPA1 female pairs showed significantly higher aggression index compared to the other five female groups. Right: aggression index in male–male assays. *d.* Pursuit index across different genotypes. Left: pursuit index in female–female assays. Right: pursuit index in male–male assays. R72A10>TRPA1 male pairs showed significantly higher pursuit index compared to the other five groups of male-male. For all six-group comparisons in *b.*, *c.*, and *d.*, significant differences were identified in all comparisons (*p* < 0.05, Kruskal–Wallis test for fly pair distance and pursuit index for female pair and male pair and aggression index for female pair, and *p* < 0.05, one-way ANOVA test for aggression index for male pair). For post-hoc pairwise testing, Dunn’s test with Bonferroni-adjusted *p*-values was used after Kruskal–Wallis test: *** *p* < 0.0005; ** *p* < 0.005; * *p* < 0.05; ns: *p* ≥ 0.05, and Tukey’s test with adjusted *p*-values was used after one-way ANOVA test: *** *p* < 0.0005; ** *p* < 0.005; ns: *p* ≥ 0.05. Pink box: experimental group of female-female. Blue box: experimental group of male-male. Gray box: control groups of the sex in comparison. Red dot: individual female-female pair. Blue dot: individual male-male pair. Brown edge border: assay group for R26E01-GAL4 labeled neurons. Orange edge border: assay group for R72A10-GAL4 labeled neurons.

To further investigate different social interactions between two flies with specific body positioning and locomotive feature, we introduced two indices - “aggression index” and “pursuit index”- by leveraging multiple body parts detected in each fly of a fly pair. (see details in “Methods”). Briefly, by using the difference between body part distances, normalized by the sum of body segment lengths of two flies, we can represent the relative body positioning of two flies. The aggression index ranges between –1 and 1, showing whether the fly pair was in “tail to tail” or “face to face” position. The pursuit index ranges between 0 and 1, showing whether the fly pair was in a random or “one’s head following the other’s tail” position, as shown in Fig. 5a. In Fig. 5.c left panel, activated R26E01>TRPA1 female-female group showed significantly higher aggression index compared to the 5 other genotype groups. Fig. 5.c right panel shows that no male-male group, including the activated R72A10>TRPA1group, displayed an elevated aggression index. The activated R72A10>TRPA1 group showed significant difference in pursuit index in the 6-genotype comparison (Fig. 5.d right panel). No group of female-female assays showed significant difference in pursuit index, including the R26E01>TRPA1 group as expected (Fig. 5.d left panel).

As shown by the aggression index and pursuit index with multiple body part**s** detection in each fly of fly pair experiment**s**, the Kestrel and multi-view integration of pose estimation models provided locomotive measurements averaged over a 30-minute recording session that were aligned with the elevated frequency of specific social interactions of a phenotype—increased aggression or courtship—beyond simple relative body distance.

### Head butt and UWE events show distinct pre-event locomotive features across different genotypes

After analyzing locomotive features over the 30-minute assay time (long timescale), we next examined individual manually scored behaviors of interest at a short timescale. We investigated the locomotive features during the 0.1 second before each tracked manually scored frame index of head butts or UWEs (see details in “Methods”). In Fig. 6.a, we analyzed the head butt events from female-female interactions across 6 different genotypes. In the left panel of Fig. 6.a, R26E01>TRPA1 female pairs’ head butts had smaller pre-event fly pair distance in comparison to the head butt events from the other 5 genotypes, which aligned with the observation of experimenters during manual scoring. Some head butts in control groups occurred in a non-reciprocal fashion when two flies were further away from each other. In the right panel of Fig. 6.a, the aggression indices just before head butts were significantly higher in the activated R26E01>TRPA1 group, consistent with the observation that head butts in this genotype typically occurred when flies were positioned head-to-head. In contrast, head butts in other genotypes occurred in more varied relative body orientations, such as one fly striking the other’s abdomen or different parts of the body.

**Figure 6.**
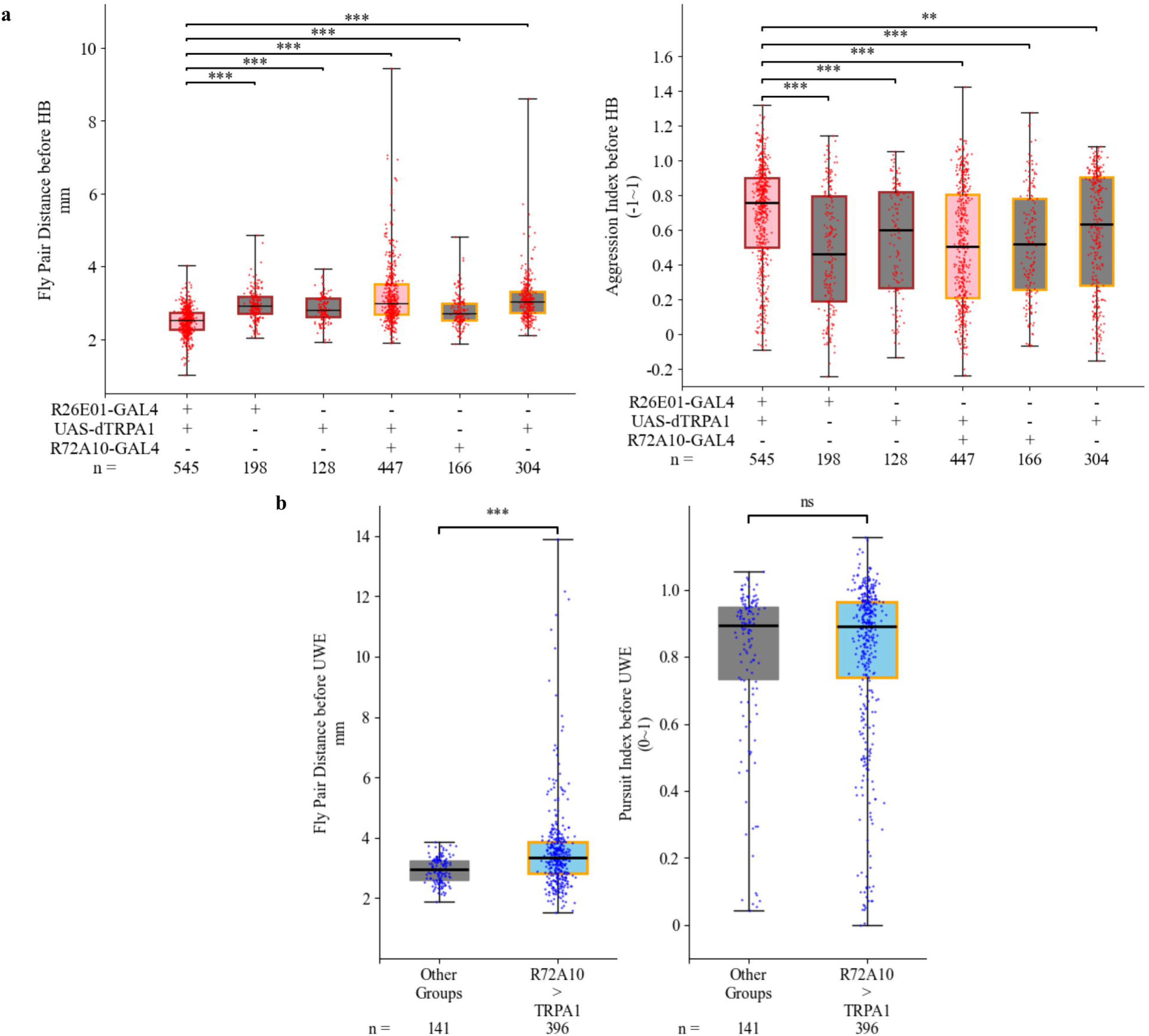
Comparison of fly pair distance, aggression index and pursuit index during 0.1 second before manual scoring event across all tracked head butt and UWE from different genotypes. *a.* Comparison of Head butt events from different genotypes. Left: fly pair distance during 0.1 second before head butt of all tracked head butts from different genotypes. Right: aggression index during 0.1 second before head butt of all tracked head butts from different genotypes. Head butt events from female-female interactions of R26E01>TRPA1 genotype were primed with shorter fly pair distance and higher aggression index between the female pair, compared to head butt events from other genotypes. Kruskal–Wallis tests were performed first, and significant differences were identified in both comparisons (*p* < 0.05, Kruskal–Wallis test). For post-hoc pairwise testing, Dunn’s test with Bonferroni-adjusted *p*-values was used: ***, *p* < 0.0005; **, *p* < 0.005; *, *p* < 0.05; ns, *p* ≥ 0.05. *b.* Comparison between UWE events from R72A10>TRPA1 line and UWE events from all the other genotypes. Due to the rarity of UWE events from male-male interactions in control groups, the comparison was performed between UWEs from R72A10>TRPA1 line, n = 396, and UWEs from all the other genotypes, n = 141. Left: fly pair distance during 0.1 second before UWE of all tracked UWEs from R72A10>TRPA1 genotype and all the other 5 genotypes. Right: pursuit index during 0.1 second before UWE of all tracked UWEs from R72A10>TRPA1 genotype and all the other 5 genotypes. UWE events from male-male interactions of R72A10>TRPA1 genotype happened in a wider fly pair distance range. Some of the UWEs happened when male-male pairs were farther apart. The pursuit index before UWE did not show a difference between two groups of UWE events. The data did not pass the test of normality by the Shapiro–Wilk normality result. Mann-Whitney U test was used for pairwise testing: ***, *p* < 0.0005; ns, *p* ≥ 0.05. Pink box: experimental group of female-female. Blue box: experimental group of male-mal e. Gray box: control groups of the sex in comparison. Red dot: individual head butt. Blue dot: individual UWE. Brown edge border: assay group for R26E01-GAL4 labeled neurons. Orange edge border: assay group for R72A10-GAL4 labeled neurons.

For individual UWE event’s locomotive feature, the comparison was done between UWEs from R72A10>TRPA1 male pairs and UWEs from the other 5 genotypes because UWEs are rare events in control male-male encounters. In Fig. 6.b left panel, the UWE events from R72A10>TRPA1 male-male interactions showed a significant difference in pre-UWE fly pair distance. In the experimental pairs, one male fly could potentially perform UWE even when two males were far away in the arena, while the fewer UWEs in the other control groups happened when two males were close in distance. Courtship songs performed by male flies in the form of UWEs to female flies have been reported to be modulated by distance between the two individuals [43,44], and the common distances discovered were similar to the male-male distances just before UWE events from this study. We speculate that the ability to perform UWEs at greater inter-fly distances may reflect the underlying mechanism by which R72A10-targeted neurons modulate courtship behavior. Additionally, in the right panel of Fig. 6.b, the body positioning just before UWE events, represented by the pursuit index before UWEs, was similar between UWEs of the R72A10>dRPA1 male pair group and UWEs of the other genotypes. This does not contradict the observation that UWEs for all the genotypes happen when one fly is chasing the other. R72A10-GAL4-targeted neurons did not affect the relative body position of two flies before performing UWEs, which usually happens while one fly is chasing another fly.

Based on locomotive feature comparison between behavior of interest from different genotypes, we validated the manual score process and discovered the subtle difference in head butts of female pair with thermally activated R26E01 neurons compared to head butts from other control groups and UWEs of male pair with thermally activated R72A10 neurons compared to UWE from other control genotypes. The kestrel experiment setup with multi-view integration of pose estimations based on neural networks could quantify fly-fly social behavior beyond the behavior count.

### R72A10-GAL4 targets ∼ 40 Neurons in the central brain including a distinct cluster at the AL-SOG junction

Finally, we sought to characterize the neurons targeted by the R72A10-GAL4 line, whose combination with TRPA1 and activation at 30 °C led to an enhanced UWE phenotype in male–male assays (Fig. 4). To the best of our knowledge, the neurons labeled by this GAL4 line have not been previously characterized. Therefore, to identify the neuronal populations targeted by R72A10-GAL4, we crossed it with the UAS-mCD8::GFP reporter and performed immunostaining using an anti-mCD8 antibody. Immunofluorescence analysis revealed that R72A10-GAL4 labeled approximately 40–45 neurons in the central brain and around 10–12 neurons (Fig. 7 a1-a3) in the ventral nerve cord (VNC) ((Fig. 7 b1-b3). In the central brain, labeled neurons were observed around the antennal lobes, the subesophageal ganglion, the lateral protocerebral complex, and several other regions.

**Figure 7.**
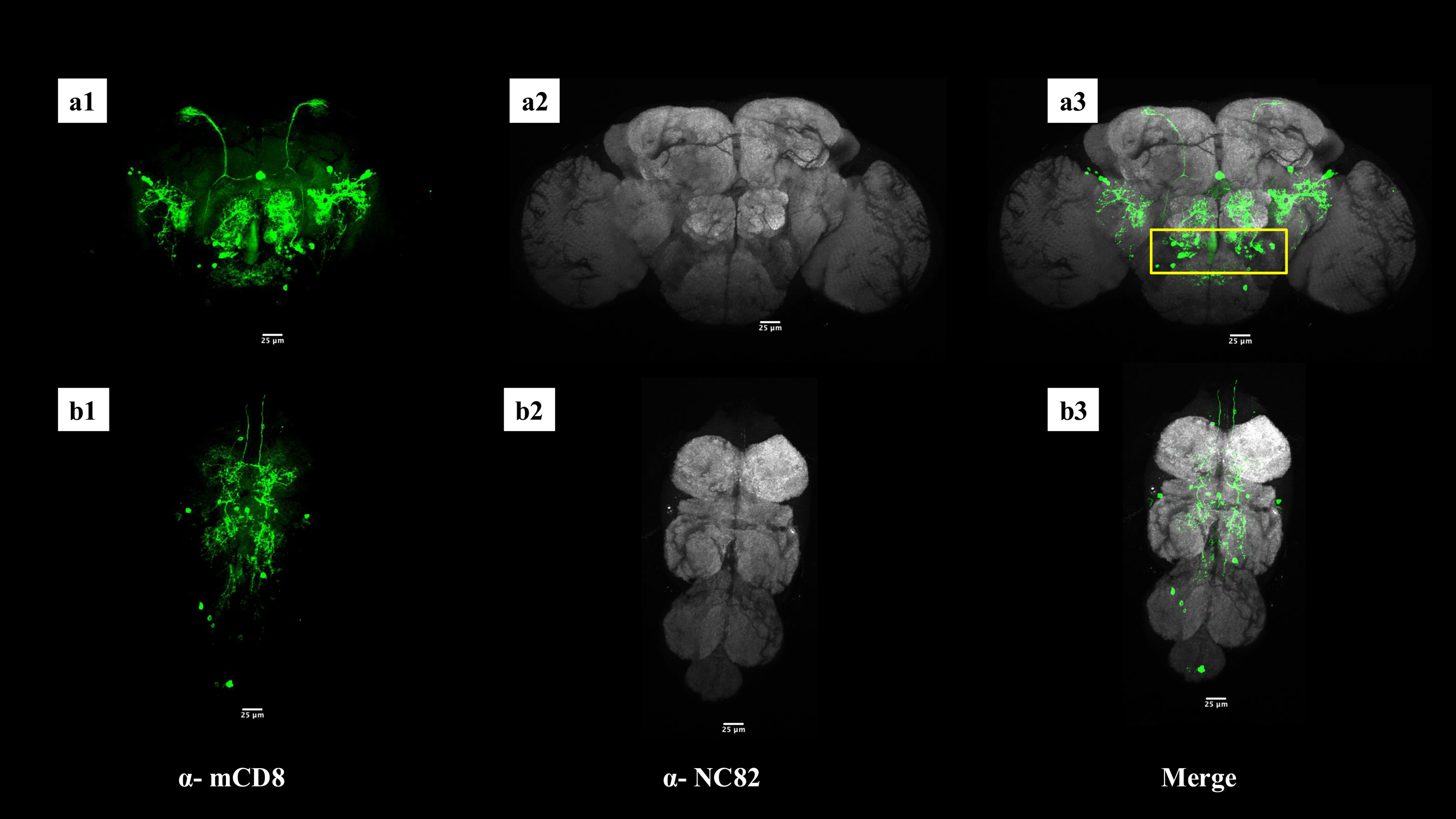
Expression pattern of R72A10-GAL4 in the adult Drosophila central nervous system. Confocal images show the expression of UAS-mCD8::GFP driven by R72A10-GAL4 in neurons of the adult central brain (a1-a3) and the ventral nerve cord (b1-b3). Neurons labeled by R72A10-GAL4 (green) are visualized using anti-mCD8 staining, while neuropil regions are labeled using anti-NC82 (gray). In the central brain, labeled neurons are located in different regions such as the antennal lobes (AL), subesophageal ganglion (SOG), and in the lateral protocerebral complex, among other regions. A distinct cluster of approximately 10–15 neurons at the junction between the AL and SOG (highlighted by yellow rectangle) is identified in each brain hemisphere. Inspection through the confocal slices reveals that this neuronal cluster has cell bodies at the AL–SOG junction (yellow rectangle) with projections extending towards higher brain regions. All images are maximum intensity projections. Scale bars, 25 µm.

Upon inspection of confocal optical sections, we identified a distinct cluster of ∼10–15 neurons located at the junction between the antennal lobes (AL) and the subesophageal ganglion (SOG) in each hemisphere of the brain (Fig. 7a3). Neuronal cell bodies in this cluster extended projections toward higher brain regions such as the lateral protocerebral complex. Given that the AL and SOG are critical sites for sensory integration of olfactory and gustatory stimuli, respectively, we hypothesized that these neurons could play a modulatory role in sensory processing circuits and, thus, behaviors^5,51^. It is plausible that this cluster contributes to the pronounced unilateral wing extension (UWE) phenotype observed in R72A10>TRPA1 males and warrants future experimental investigation.

Given the established role of octopamine in modulating male behavioral choices in *Drosophila*, we next investigated whether this cluster was octopaminergic^51–53^. To determine this, we performed double immunostaining on male brains from R72A10-GAL4/UAS-mCD8::GFP flies using antibodies against TDC2 (anti-Tdc2), an enzyme essential for octopamine synthesis^51^, and mCD8. Our results revealed no co-labeling of these neurons with anti-TDC2, indicating that they are not octopaminergic (Supplementary figure 1). At present, the neurotransmitter identity of this neuronal cluster remains unknown. Though not octopaminergic themselves, it is also that these neurons may interact with octopaminergic circuits, potentially modulating sensory integration pathways that ultimately influence male behavioral outcomes, such as the elevated UWE phenotype observed in male–male encounters.

## Discussion

Combining state-of-the-art imaging systems and deep learning-based animal pose estimation solutions, we have established a novel pipeline for investigating social behaviors in dyadic fruit fly assay and quantified them via locomotive features across both long and short timescales. The custom Kestrel imaging system, powered by Ramona’s MCAM technology, achieved high throughput fly pair behavioral assay with high temporal and spatial resolution video recording. A 3D-printed modular chamber array provides an efficient way to prepare multiple behavioral experiments with minimal stress during fly transfer, while offering ample 3D space for free movement of each fly during the assay. Multi-view integration of video inferencing results from 3 DeepLabCut models based on different body orientations of a fly in single camera view was developed as open-source approach in Python environment, which provided comprehensive body part coordinate representation during the whole recording session at 60 Hz, with minimal no-detection gaps

To evaluate the system’s reproducibility, effectiveness, and efficiency in detecting social phenotypes, we tested two GAL4 driver lines—R26E01 and R72A10—crossed to the thermogenetic effector line TRPA1. Same-sex, age-matched flies reared in isolation were recorded under heat-activated conditions. Across all recordings, fly pair detection rate exceeded 92% at 60 Hz and 95% at 10 Hz. To further assess fidelity of fly pair detection in studying behaviors, we defined an “event tracking rate” as a criterion to see if the pose estimation implementation was selectively missing body part tracking at the manually scored timestamps of behaviors of interest, and despite higher requirement threshold, event tracking rates of head butt, UWE and lunge were comparable to fly pair detection rate, at around 90%. This multi-view approach mitigates common pose estimation challenges, including those arising from unrestrained fly movement, occlusion, and inconsistent tracking quality.

Using female and male pairs from a control group (TRPA1×w^1118^), we evaluated the pose estimation results based on known features of fruit fly anatomy and locomotion from previous studies and our own observations in this study. The system was able to measure whole body length and body segment length of individual flies, and the results aligned with the known differences in female and male body size^47^ and could be used as a screening tool to confirm the sex of subjects in a fly experiment when social interaction is absent. Locomotive features were compared between female pairs and male pairs. We found that there was no significant difference in average fly pair distance between female pairs and male pairs. Consistent with a previous study^48^, male–male pairs, after isolation in confined space, exhibited increased movement; this was reflected in their higher fly pair traversal speed compared to their female counterparts. Manual counts of aggressive behaviors—head butts in female pairs and lunges in male pairs—were compared against locomotive features. The linear correlations between behavior counts and locomotive metrics across the experimental duration did not contradict the observed social interactions in control groups.

An important aspect of the system was its capacity for high-throughput behavior recording that enabled clear phenotype observation alongside moment-to-moment locomotive features. To assess whether our setup could faithfully replicate previously published results, we conducted dyadic fruit fly behavioral assays in which neurons targeted by the R26E01-GAL4 driver were thermogenetically activated using the TRPA1 thermosensitive cation channel at 30C. In line with prior findings ^9^, manual annotation of Kestrel-recorded videos revealed elevated head butt frequency in R26E01>TRPA1 female pairs, consistent with an aggression-promoting effect. In contrast, no significant increase in aggression was observed in male pairs of the same genotype. Next, we activated a distinct population of approximately 40 neurons targeted by R72A10-GAL4, a previously uncharacterized driver, and manually annotated behaviors from Kestrel-recorded dyadic assays. In male–male assays, thermogenetic activation of R72A10 neurons resulted in a robust increase in unilateral wing extension (UWE), a well-established marker of courtship behavior. This increase occurred without any detectable change in aggression. R72A10>TRPA1 females on the other hand did not show increased aggression or courtship behavior upon activation. Control male–male assays were dominated by aggressive interactions with minimal or no courtship. Therefore, the observed enhancement of male–male courtship upon activation of R72A10 neurons suggests that this neuronal population shifts behavioral output toward courtship in this social context.

To probe the neural basis of the R72A10-driven behavioral phenotype, we examined the anatomy and neurochemical identity of the labeled neurons. Among ∼40–45 central brain neurons targeted by the R72A10-GAL4, a distinct cluster of ∼10–15 was located at the AL–SOG junction, a region involved in integrating olfactory and gustatory cues. These neurons projected to higher-order centers, suggesting a role in modulating sensory-driven social behavior. To further explore the neurochemical identity of this cluster, we performed immunostaining for octopaminergic markers, given the well-established role of octopamine in regulating male social behavior in *Drosophila*. The neurons at the AL–SOG junction were not octopaminergic, and their transmitter identity remains unknown. Further immunostaining for additional neurotransmitter markers and genetic assays will be necessary to resolve their molecular profile.

In this study, we used fly pair distance and traversal speed as locomotive indicators of increased aggression (enhanced head butts) or courtship (enhanced UWEs). Intuitively, fly pair distance reflects the frequency or intensity of social interaction, and our system reliably detected significant differences in this measure over a 30-minute window between female pairs with high head butt counts or male pairs with high UWE counts, compared to their respective controls. In contrast, fly pair traversal speed over 30 minutes was not a statistically significant predictor of social behavior.

By leveraging multi–body part detection, we created an aggression index and a pursuit index—relative body position measures—to represent different types of fly pair social interaction, namely aggression and courtship, during a recording session by MCAM. By combining the two experimental sets, we tested one experimental group against five control groups, enabling a stricter statistical comparison than a standard three-group analysis. The aggression index strongly indicated elevated aggression in the R26E01>TRPA1 female pairs, whereas R72A10>TRPA1 male pairs with frequent UWEs did not differ significantly in aggression index compared to the control groups. Conversely, male pairs from the R72A10>TRPA1 group with frequent UWEs showed a significantly higher pursuit index than control male pairs. These locomotive features, derived from pose estimation, demonstrate that the Kestrel system supports high-throughput, high-resolution recording of fly social interaction, and that post-experiment body part tracking is sufficient to capture consistent interaction phenotypes over multi-minute timescales.

Furthermore, we investigated locomotive features on the timescale of individual events—head butts and UWEs—in fly pairs of different genotypes. For the R26E01 driver line, in addition to increased aggression as indicated by higher head butt counts in female pairs, we found that fly pair distance immediately before head butts was shorter in the experimental group than in the five control groups. Aggression index prior to head butts was also higher in the experimental group, suggesting that head butts occurred more frequently in a reciprocal rather than one-sided manner. Pursuit index before UWEs in R72A10>TRPA1 male pairs was not significantly different from that of control male pairs, consistent with the observation that UWE—though rare in control males—often occurred when one fly was chasing another, producing a similar body position profile as that observed in R72A10-activated flies. However, UWE events in the experimental group more frequently occurred at greater fly pair distances, whereas in control groups, UWEs were almost exclusively observed when the flies were in proximity. These differences in locomotive features preceding well-defined behaviors such as head butts and UWEs may reflect distinct motivational states, modulated by different neural or circuit-level mechanisms.

Additional behavioral data, baseline recordings, and refined feature extraction will be essential for developing closed-loop fly social behavior systems capable of early behavior prediction and real-time stimulation control^54^. This study demonstrates that high-resolution, high-throughput behavioral tracking, combined with multi-body part pose estimation, enables precise quantification of social behaviors in *Drosophila* over extended timescales. By capturing both fine-scale locomotor dynamics and discrete behavioral events such as head butts and unilateral wing extensions, the MCAM-based pipeline bridges traditional manual scoring with scalable computational approaches. The integration of spatial tracking, pose-derived behavioral indices, and genotype-specific manipulation revealed how distinct neural populations modulate aggression and courtship in sex- and context-dependent ways. Notably, the system detected subtle but behaviorally meaningful variations—such as differences in pairwise distance and pursuit—both across entire assays and immediately preceding individual events. These features not only validated established phenotypes but also uncovered new spatiotemporal signatures that may reflect underlying motivational states. As a flexible, open-source platform, this pipeline offers a robust tool for linking neural circuit activity to behavior and provides a foundation for future closed-loop experiments and predictive modeling in freely moving animals.

## Methods

### Fly Stocks

Flies were reared on standard cornmeal-sucrose-yeast-agar food under 50% relative humidity and maintained on a 12 h light / 12 h dark cycle. The following *Drosophila melanogaster* stocks were obtained from the Bloomington Drosophila Stock Center: R26E01-GAL4 (#60510), R72A10-GAL4 (#48306), UAS-mCD8::GFP (#32185), and UAS-dTRPA1 (#26264).

### Fly Rearing and Preparation for Behavioral Experiments

All fly crosses were performed at 19 °C. During the late pupal stage, individual flies were transferred to glass vials (10 mm × 75 mm; VWR catalog #47729-568) containing 1 mL of standard cornmeal-sucrose-yeast-agar food. Flies eclosed in these vials, remained socially isolated throughout adulthood, and subsequently assayed for behaviors at 30 °C for TRPA1 thermogenic activation.

Shortly after eclosion, each fly was briefly anesthetized with CO and marked on the dorsal thorax with a small paint dot. To enable individual identification during experiments and data recording, two paint colors were used: one fly received a blue paint dot and the other received a white paint dot. Flies were returned to their original vials for recovery and allowed at least 48 hours before behavioral testing. Behavioral experiments were performed using same-sex, same-genotype pairings of age-matched flies (10-24 days old).

### Behavioral Assay Setup and Recording

Fly behavioral assay was conducted using the Kestrel, which enables simultaneous high-resolution behavioral recording across multiple arenas with a wide field of view and high temporal resolution. A custom-designed Kestrel-compatible 2x4 arena chamber array was used to conduct eight parallel same-sex fly-fly interactions per recording session. The floor of each chamber contained a central 7 mm circular food patch consisting of 70 µL standard fly food topped with 10 µL yeast solution to promote fly interactions. Each chamber was equipped with a movable divider. Via the opening on each side of the chamber, each fly was transferred via phototaxis and gravitaxis from individual glass vial into the chamber without stress of air aspiration or other physical contact, and two flies for social interaction experiment were kept separated before recording started.

At the start of each recording session, the eight arena chambers were slotted into fixed positions on the Kestrel arena base plate. Dividers were retracted simultaneously with the onset of video acquisition, allowing paired flies to interact freely. Each session lasted 35 minutes, and the initial 30-minute encounter was used for manual scoring and automated data analysis. The Kestrel system automatically parsed the composite video into eight individual fly-fly recordings (one per chamber) at 60 Hz and 576 × 576-pixel resolution. Additional details on the arena design, chamber configuration, and the Kestrel imaging setup are provided in Fig. 1 of the Results section.

### Computing Hardware and Software

Two desktop computers running Ubuntu operating systems were used in the study for computing purposes. One desktop computer in lab space with Intel Xeon(R) W-2245 and Quadro RTX 5000 on Ubuntu 22.04.3 LTS was used as part of the Kestrel to record fly pair assay and run DLC inference. The other desktop computer in office space with AMD Ryzen 7 7700x and NVIDIA GeForce RTX 4070 Ti SUPER on Ubuntu 24.04.01 LTS was used for the purposes of general coding, DLC model training, and DLC inference. All analyses and algorithms were conducted in Python 3.10 using Jupyter Notebook for code execution and data visualization.

### DLC Implementation

#### DLC model training

We used DLC v3.0.0 for pose estimation. Specifically, we used a ResNet-152-based neural network with default parameters, validated with a 95% training and 5% testing split and 1 shuffle. For the top view model, we labeled 305 frames from clips of manually scored head butt, lunge, and UWE events when fly pairs were in the top view angle, with 12 body parts, including head, left eye, right eye, center, tail, wing close, left wing point 0, left wing point 1, left wing point 2, right wing point 0, right wing point 1, and right wing point 2. After 100000 training iterations, the top view model had a training error of 2.66 pixels and a test error of 3.26 pixels. For the side view model, we labeled 300 frames from clips of manually scored head butt, lunge, and UWE events when fly pairs were in the side view angle, with 7 body parts, including mouth, head, center, tail, wing close, wing extend 0, and wing extend 1. After 90000 training iterations, the side view model had a training error of 3.14 pixels and a test error of 3.49 pixels. 158 frames from the top view and side view model inference result were used to train the cross view model, with 8 body parts, including “top”, “top left”, “top right”, “left”, “right”, “down left”, “down right” and “down”. After 30000 training iterations, the cross view model had a training error of 3.36 pixels and a test error of 4.40 pixels. Body parts such as head, center, and tail from the top and side view models and “top”, “left”, “right”, and “down” from the cross view model were used for locomotion analysis. All the training clips for the top view and side view model training were from 18 fly-fly videos from the first 4 experiment recording days for R72A10-GAL4 labeled neuron activation. Outlier frames used for cross view model training were taken from 5 fly-fly videos recorded on the first experiment day.

#### Pre-DLC-inference Preparations

The MCAM software automatically crops out fly-fly video of 576 by 576 pixels based on the custom chamber after a recording session. However, further cropping was performed to recenter the arena in each video due to slight differences in chamber location in the camera view and to accelerate the DLC inference process. Using HoughCircles() in the opencv2 Python module, the center and inner radius of the arena were determined in pixel units. The inner radius of the arena in pixel units across all videos ranged between 225 to 235 for a physical size of 8.4 mm, so a pixel-to-physical scale ratio was calculated for each fly-fly video. During DLC inference, further cropping was performed based on the center location of the arena and extending 240 pixels in all 4 directions, which covers the entire arena inner radius.

### Fly Body Segment Length Measurement

The body segment length of individual fly was measured by using a section of video recording with fly pair pose estimation results of high confidence from top or side view model. The section of pose estimation result is an at least 1-second-long time window meeting three conditions: (1) All head, center, and tail are detected from both flies at every frame in the time window; (2) The head-to-head, tail-to-tail, and head-to-tail distances between the fly pair are at least 1mm apart to guarantee clear separation of two flies in view; (3) With 6mm distance between center body part to arena center as threshold, it is decided whether the section is from top or side view model. If both center body parts are within the 6mm threshold, then both flies are on the floor in top view angle and, hence, the section for body segment length measurement is from top view model pose estimation result. If both center body parts are outside the 6mm threshold, the side view model pose estimation is used.

The average distance between the front part, f, and the rear part, r, is defined as the length of the body segment, d(f, r). The front and rear part are head and tail for whole body segment, head and center for head+thorax segment, or center and tail for abdomen segment.

### View-Selection Filter for Multi-View Integration

The view-selection filter is based on DLC pose estimations’ confidence number and fly realistic physical size. Following priorities steps and rules, the filter fills the gap of the head, center, and tail detections of fly pair in the top view model baseline to form multi-view coordinate array (MVCA).

In step 1, the confidence number (conf: 0 to 1) from the DLC output is used to decide which pose estimation from the top view model is used directly. From top view model inference, pose pairs with conf > 0.95 are used in the MVCA. For example, fly 1 tail with conf = 1 and fly 2 tail with conf = 0.99, then the tail pair estimation is defined as tail pair locations in the MVCA.

Step 2 utilizes frames when only one specific body part of a single fly is detected in the top view model. The other fly’s said body part is searched for in pose estimation from side view model inference. Three thresholds are used to decide if the side view body part is to be paired with the top view counterpart: (1) conf is > 0.99; (2) distance threshold from top view counterpart is >0.5mm (∼15 pixels); (3) the one of two from side view is further away from the fly with top view counterpart.

After utilizing all the viable detection from top view, step 1 and step 2 were repeated for all the remaining frames with side view model inference to fill the remaining undefined pose pair in MVCA. The only difference here is an increased distance threshold, 3mm (∼100 pixels), significantly larger than the length of an average fly. At this step, single-fly body part pose estimation, not pair, from the side view model is still being considered because the top view model did not provide any pose estimation about this specific body part. So this specific body part pose estimation’s viability is decided by its relative location to known pose estimations of other body parts in MVCA. So a larger distance threshold is used for a conservative approach.

When all the head, center and tail pose estimations from top and side view models are depleted, cross view model inference is used to fill the gap in fly pair observation in the final step. 4 poses in cross view model have a strict DLC conf threshold of 1.0, and “cross”, the average of 4 poses, is used as the single body part in the cross view. For timestamps without pose estimation of either fly in MVCA, viable “cross” pair in cross view model fills in MVCA. For frames with pose estimation of single fly in MVCA, “cross” coordinate that is farther away from the known fly location in MVCA will be used in MVCA for the other fly.

Besides fly pair MVCA at 60 Hz from original 60fps video recording, down-sampled results at 10 Hz and 1 Hz are also used, by averaging tracked body part coordinates in respective sampling windows. The threshold for if a body part is tracked at down-sampled timestamp is 10% of the 60 Hz data points within the lower rate window. For example, 10 Hz sampling window length is 6, so as long as there is one frame that returns viable body part coordinate in the sampling window, the timestamp for 10 Hz sampling rate is defined as viable. For the 1 Hz sampling window, the threshold is 6 data points out of a 60-frame window.

### Fly Pair Tracking Performance Evaluation

#### Fly Pair Detection Rate

Fly pair detection rate (%) is defined to evaluate how completely the integrated body part tracking result covers the fly pair’s location in the custom arena during the experiment time. To achieve an accurate representation of the body positioning of two flies, at least two pairs of body parts from both flies are required to be tracked when both flies are in the top view or side view angle. For example, if the body part tracking result at one timestamp has one fly’s center and tail coordinates, and the other fly’s head, center, and tail coordinates, this frame is defined as a viable fly pair detection because both flies have center and tail coordinates estimated. For another example, if one fly has head and center coordinates but the other has center and tail coordinates, this frame is defined as a nonviable fly pair detection. This criterion reduces inaccuracy due to misidentification of the owner of the body part. Despite extensive filtering, as in the example, we can only have very high confidence that the two detected center body parts are from two different flies, and lower confidence that the head and tail are correctly assigned to different flies. Another case of viable fly pair detection occurs when both flies are in the cross view angle, or when one fly is in the top or side view angle and the other is in the cross view angle. The f fly in the floor or side view angle is required to have at least two body part detections. The definition of fly pair detection rate provides a baseline for other locomotion parameters used in the study. Fly pair detection rate also represents the percentage of time that the following locomotion parameters are viable during the experiment.

#### Event Tracking Rate

Built upon the fly pair detection rate, event tracking rate (%) is a metric that quantifies the proportion of successfully tracked behavioral events relative to the total number of manually counted events. A successfully tracked event is defined when the event window of 5 frames (1/12 of a second), centered around the frame timestamp of the manually scored event, has a fly pair detection rate equal to or higher than 80% - that is, at least 4 out of 5 frames have viable fly pair detection.

### Locomotion Parameters

#### Fly Pair Distance

Fly pair distance (mm) is the Euclidean distance between two flies’ body centers. There are 3 ways to acquire approximate body center locations: (1) the average of head and tail coordinates; (2) the coordinate of center body part; (3) the cross coordinate when a fly is in cross view angle. The body center of one fly is defined as an average of viable approximate body center locations from a frame.

#### Fly Pair Traversal Speed

Fly pair traversal speed (mm/s) is used to define how much distance a fly pair as whole traverses inside a custom arena during a specific period, an average of two flies’ body center speed. To focus on measuring traversal distance p not the localized bursts of movements, exemplified by sustained fencing, female head butt or male lunge, multi-view coordinate array at 10 Hz sampling rate is used to calculate the speeds of two tracked flies, 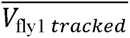 and 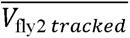 as average body center displacement between frames divided by 0.1 second due to 10 Hz sampling rate.

To reduce the error of tracking identification switching during the whole experiment, fly pair traversal speed, 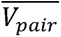, is defined to reliably estimate as shown in the equation below. In this example case, if there was a misidentification for 5% of whole tracking period, the average result of speeds of two tracked flies is approximately equal to average of speeds of two flies with true identifications, 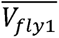 and 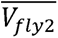:

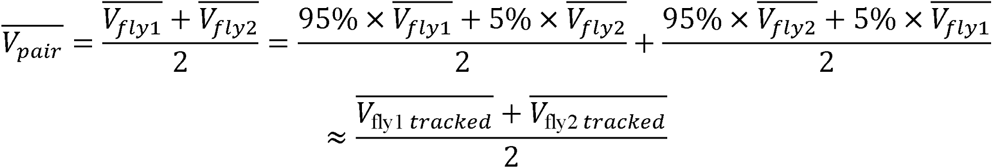

This calculation removed the segments of potential misidentification during the tracking period, and only timestamps of identity switch were not considered.

#### Aggression Index and Pursuit Index

To depict social behavior of two flies, aggression index and pursuit index are introduced as normalized position parameters utilizing at least two pairs of body part detections from head, center, and tail of top view and side view models’ combination. As shown in Fig. 5.a, two pairs of body parts from two flies includes front part of fly 1, f1, rear part of fly 1, r1, front part of fly 2, f2, and rear part of fly 2, r2. The summation of body segment measurements of the two flies:

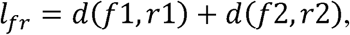

was used as a divisor in the normalization process for each fly pair detection situation.

Specifically, aggression index measures if the flies are “front to front” or “rear to rear” by using distance between two flies rear parts, *d*(*r*1, *r*2), and distance between two flies front parts, *d*(*f*1, *f*2):

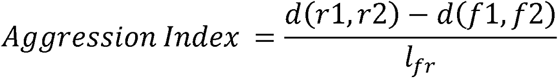

Because the theoretical difference between rear parts distance and front parts distance is between -*l_fr_* and *l_fr_*, the aggression index is theoretically a value between –1 and 1 after normalization, with a higher value indicating two flies are in a relative position of “front to front”.

For pursuit index, it measures if the flies are “front to rear” or not “front to rear” by using distances between front part of one fly and rear part of the other fly, d(f1, r2) and *d*(*f*2, *r*1):

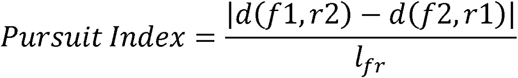

Because the theoretical absolute difference between distances between fly’s front part to the other fly’s rear part is between 0 and *l_fr_*, the pursuit index is theoretically a value between 0 and 1, with a higher value indicating that two flies are in a relative position of one chasing or following the other.

Fly pair distance, aggression index and pursuit index before an event was calculated for all tracked HB and UWE events. Based on the criterion for tracked event, every tracked event has viable fly pair detection in 0.1-second time window of 6 frames, including the frame of manually scored timestamp of the event and 5 frames preceding it. Hence, the average value of locomotion parameters in the 0.1-second time window, including fly pair distance, aggression index and pursuit index is defined as “locomotion parameter before event”, for instance, “aggression index before HB” or “fly pair distance before UWE”. Due to pose estimation error at each frame, these average aggression and pursuit indexes over a short time window may fall outside the theoretical value range.

### Statistics Analysis

For two-group comparisons, the Shapiro-Wilk test of normality was used to initially determine data distribution for each group. If p > 0.05, the data group was determined to have a normal distribution. The Mann-Whitney U test was used for two groups that were non-normally distributed, and the independent t-test was used for two groups that were both normally distributed. For correlation analysis between a pair of data points, Pearson’s R² and p-value were used. For comparisons of three groups and six groups in this study, the Shapiro-Wilk test of normality was performed for each group of data points. A one-way ANOVA test was used for normally distributed groups, and Tukey’s test with adjusted p-values was then performed as a post hoc pairwise comparison. For non-normally distributed groups, the Kruskal-Wallis test was performed, and for tests with p values < 0.05, significant differences were identified among the groups. For post hoc pairwise comparisons, Dunn’s test with Bonferroni-adjusted p-values was performed. All box plots in the study were plotted with the first and third quartiles as the box boundaries, the median line in the box, and whiskers extending to the minimum and maximum values of the data groups. For a detailed list of statistical tests performed and associated p-values, see Supplementary Data 2: *Statistics*.

### Immunostaining and Imaging

Adult male brains and ventral nerve cords (VNC) were dissected, fixed, and stained as described previously^42,49^. The following primary antibodies were used: rat α-mouse CD8 (1:100, Invitrogen, MCD0800), mouse α-nc82 (1:20, Developmental Studies Hybridoma Bank), and rabbit α-TDC2 (1:100, Covalab). Secondary antibodies conjugated to Alexa Fluor 488, Alexa Fluor 594, or Alexa Fluor 647 (Molecular Probes) were used at a dilution of 1:200. Labeled brains were mounted in Vectashield (Vector Labs, #H1000). All images are confocal serial sections acquired using an Olympus FluoView FV1000 microscope with either a 20×/NA 0.75 (air) objective or a 60×/NA 1.42 (oil) objective and processed in Fiji (https://fiji.sc/). Maximum intensity projections were generated for visualizing the neurons.

## Supporting information

Supplementary Data 1: Event Tracking Rate

Supplementary Data 2: Statistics

## Code availability

All jupyter notebook and python scripts used in the study is at the link: https://github.com/ziyingc/flypairtracking

## Author Contributions

C.B.P.-M. and S.S. conceived the project. C.B.P.-M., S.S., and Z.C. contributed to experimental design, data interpretation, and manuscript development. S.S. and Z.C. agreed that either author can be listed first in reporting the manuscript. J.E., A.B., and M.H. developed the hardware and software and built the Kestrel system in close collaboration with S.S., who provided biological and behavioral design input, standardized aggression assays, and contributed to system testing and refinement. Z.C. performed the majority of behavioral experiments and independently wrote the code for downstream processing and analysis of Kestrel-recorded videos. A.S.A., A.E.U., and S.Y. contributed to additional behavioral assays. Y.B.C. carried out the immunostaining experiments. S.A.H. analyzed immunostaining images using FIJI and generated maximum intensity projection data. The manuscript was written, reviewed, and edited by C.B.P.-M., S.S., and Z.C., with feedback from A.B., Y.B.C., and S.Y.

## Acknowledgement

We are grateful to Kay Bhagabani for reviewing the manuscript and providing helpful comments.

This work was supported by institutional funds from McLean Hospital Corporation and by the National Institute of General Medical Sciences of the National Institutes of Health under award numbers **5K99GM141449**, **3R00GM141449-05S1**, and **3R00GM141449-03S1**, awarded to **Caroline B. Palavicino-Maggio**. Additional support was provided by the Office of the Director, National Institutes of Health, under award number **R44OD036187**, awarded to **Ramona Optics Inc.**

---

**Supplementary Video 1 |**

Representative video of two R26E01>TRPA1 female flies during thermogenetic activation. The flies exhibit elevated headbutt frequency, as previously reported^9^. Colored dots denote tracked fly centroids. The number at the upper left indicates fly pair distance (in mm), while the number at the lower left shows the frame index of the experiment from which this behavioral sequence was extracted.

**Supplementary Video 2 |**

Representative video of a dyadic encounter between two R72A10>TRPA1 male flies during thermogenetic activation. The males display elevated unilateral wing extensions (UWEs), a behavior that is rarely observed in control male–male encounters. Colored dots indicate tracked fly centroids. The number at the upper left indicates fly pair distance (in mm), while the number at the lower left shows the frame index of the experiment from which this behavioral sequence was extracted.

**Supplementary Figure 1 |**

Confocal images showing higher magnification views of the R72A10-GAL4-labeled neuronal cluster at the AL–SOG region in a male adult Drosophila brain. The neuronal cluster (green, left) at the AL–SOG junction does not show co-localization with anti-Tdc2-positive neurons (magenta, middle) in the merged image (right), indicating that this cluster is not octopaminergic. Scale bars, 25 µm

## Bibliography

1. Fernández, M. P. & Kravitz, E. A. Aggression and courtship in Drosophila: pheromonal communication and sex recognition. J. Comp. Physiol. A Neuroethol. Sens. Neural Behav. Physiol. 199, 1065–1076 (2013).

2. Sato, K., Goto, J. & Yamamoto, D. Sex mysteries of the fly courtship master regulator fruitless. Front. Behav. Neurosci. 13, 245 (2019).

3. Marie-Orleach, L., Bailey, N. W. & Ritchie, M. G. Social effects on fruit fly courtship song. Ecol. Evol. 9, 410–416 (2019).

4. Asahina, K. Neuromodulation and strategic action choice in Drosophila aggression. Annu. Rev. Neurosci. 40, 51–75 (2017).

5. Wang, L. & Anderson, D. J. Identification of an aggression-promoting pheromone and its receptor neurons in Drosophila. Nature 463, 227–231 (2010).

6. Kravitz, E. A. & Fernandez, M. de la P. Aggression in Drosophila. Behav. Neurosci. 129, 549–563 (2015).

7. Chen, S., Lee, A. Y., Bowens, N. M., Huber, R. & Kravitz, E. A. Fighting fruit flies: a model system for the study of aggression. Proc. Natl. Acad. Sci. U. S. A. 99, 5664–5668 (2002).

8. Chiu, H., et al. A circuit logic for sexually shared and dimorphic aggressive behaviors in Drosophila. Cell 184, 847 (2021).

9. Palavicino-Maggio, C. B., Chan, Y.-B., McKellar, C. & Kravitz, E. A. A small number of cholinergic neurons mediate hyperaggression in female Drosophila. Proc. Natl. Acad. Sci. U. S. A. 116, 17029–17038 (2019).

10. Asahina, K., et al. Tachykinin-expressing neurons control male-specific aggressive arousal in Drosophila. Cell 156, 221–235 (2014).

11. Hoopfer, E. D. Neural control of aggression in Drosophila. Curr. Opin. Neurobiol. 38, 109–118 (2016).

12. Watanabe, K., et al. A circuit node that integrates convergent input from neuromodulatory and social behavior-promoting neurons to control aggression in Drosophila. Neuron 95, 1112–1128.e7 (2017).

13. Nilsen, S. P., Chan, Y.-B., Huber, R. & Kravitz, E. A. Gender-selective patterns of aggressive behavior in Drosophila melanogaster. Proc. Natl. Acad. Sci. U. S. A. 101, 12342–12347 (2004).

14. Fan, P., et al. Genetic and neural mechanisms that inhibit Drosophila from mating with other species. Cell 154, 89–102 (2013).

15. Ahrens, M. B., Huang, K. H., Narayan, S., Mensh, B. D. & Engert, F. Two-photon calcium imaging during fictive navigation in virtual environments. Front. Neural Circuits 7, 104 (2013).

16. Harvey, C. D. & Svoboda, K. Locally dynamic synaptic learning rules in pyramidal neuron dendrites. Nature 450, 1195–1200 (2007).

17. Seelig, J. D., et al. Two-photon calcium imaging from head-fixed Drosophila during optomotor walking behavior. Nat. Methods 7, 535–540 (2010).

18. Kohatsu, S. & Yamamoto, D. Visually induced initiation of Drosophila innate courtship-like following pursuit is mediated by central excitatory state. Nat. Commun. 6, 6457 (2015).

19. Thomson, E. E., et al. Gigapixel imaging with a novel multi-camera array microscope. Elife 11, (2022).

20. Johnson, R. E., et al. Probabilistic models of larval zebrafish behavior reveal structure on many scales. Curr. Biol. 30, 70–82.e4 (2020).

21. Reiter, S., et al. Elucidating the control and development of skin patterning in cuttlefish. Nature 562, 361– 366 (2018).

22. Buchanan, S. M., Kain, J. S. & de Bivort, B. L. Neuronal control of locomotor handedness in Drosophila. Proc. Natl. Acad. Sci. U. S. A. 112, 6700–6705 (2015).

23. Lohmann, A. W. Scaling laws for lens systems. Appl. Opt. 28, 4996–4998 (1989).

24. Chowdhury, B., Wang, M., Gnerer, J. P. & Dierick, H. A. The Divider Assay is a high-throughput pipeline for aggression analysis in Drosophila. *Commun*. Biol. 4, 85 (2021).

25. Yadav, R. S. P. et al. DANCE: An open-source analysis pipeline and low-cost hardware to quantify aggression and courtship in Drosophila. bioRxiv (2025) doi:10.1101/2025.01.03.631168.

26. 26. Eyjolfsdottir, E. et al. Detecting social actions of fruit flies. in Lecture Notes in Computer Science 772–787 (Springer International Publishing, Cham, 2014).

27. Kabra, M., Robie, A. A., Rivera-Alba, M., Branson, S. & Branson, K. JAABA: interactive machine learning for automatic annotation of animal behavior. Nat. Methods 10, 64–67 (2013).

28. Mathis, A., et al. DeepLabCut: markerless pose estimation of user-defined body parts with deep learning. Nat. Neurosci. 21, 1281–1289 (2018).

29. 29. Nath, T., et al. Using DeepLabCut for 3D markerless pose estimation across species and behaviors. bioRxiv (2018) doi:10.1101/476531.

30. Lauer, J., et al. Multi-animal pose estimation, identification and tracking with DeepLabCut. Nat. Methods 19, 496–504 (2022).

31. Pereira, T. D., et al. SLEAP: A deep learning system for multi-animal pose tracking. Nat. Methods 19, 486– 495 (2022).

32. McKenzie-Smith, G. C., Wolf, S. W., Ayroles, J. F. & Shaevitz, J. W. Capturing continuous, long timescale behavioral changes in Drosophila melanogaster postural data. PLoS Comput. Biol. 21, e1012753 (2025).

33. Klibaite, U., Berman, G. J., Cande, J., Stern, D. L. & Shaevitz, J. W. An unsupervised method for quantifying the behavior of paired animals. Phys. Biol. 14, 015006 (2017).

34. Potsaid, B., Bellouard, Y. & Wen, J. Adaptive Scanning Optical Microscope (ASOM): A multidisciplinary optical microscope design for large field of view and high resolution imaging. Opt. Express 13, 6504–6518 (2005).

35. Ashraf, M., et al. Random access parallel microscopy. Elife 10, e56426 (2021).

36. Orth, A. & Crozier, K. Gigapixel fluorescence microscopy with a water immersion microlens array. Opt. Express 21, 2361–2368 (2013).

37. Duffy, J. B. GAL4 system in Drosophila: a fly geneticist’s Swiss army knife. Genesis 34, 1–15 (2002).

38. Jones, W. D. The expanding reach of the GAL4/UAS system into the behavioral neurobiology of Drosophila. BMB Rep. 42, 705–712 (2009).

39. Kallman, B. R., Kim, H. & Scott, K. Excitation and inhibition onto central courtship neurons biases Drosophila mate choice. Elife 4, e11188 (2015).

40. Trannoy, S., Chowdhury, B. & Kravitz, E. Handling alters aggression and “loser” effect formation in Drosophila melanogaster. Learn. Mem. 22, 64–68 (2015).

41. Barbagallo, B. & Garrity, P. A. Temperature sensation in Drosophila. Curr. Opin. Neurobiol. 34, 8–13 (2015).

42. Alekseyenko, O. V., et al. Single serotonergic neurons that modulate aggression in Drosophila. Curr. Biol. 24, 2700–2707 (2014).

43. Clowney, E. J., Iguchi, S., Bussell, J. J., Scheer, E. & Ruta, V. Multimodal chemosensory circuits controlling male courtship in Drosophila. Neuron 87, 1036–1049 (2015).

44. Koganezawa, M., Kimura, K.-I. & Yamamoto, D. The neural circuitry that functions as a switch for courtship versus aggression in Drosophila males. Curr. Biol. 26, 1395–1403 (2016).

45. Yamamoto, D. & Koganezawa, M. Genes and circuits of courtship behaviour in Drosophila males. Nat. Rev. Neurosci. 14, 681–692 (2013).

46. Schretter, C. E., et al. Cell types and neuronal circuitry underlying female aggression in Drosophila. Elife 9, (2020).

47. Testa, N. D., Ghosh, S. M. & Shingleton, A. W. Sex-specific weight loss mediates sexual size dimorphism in Drosophila melanogaster. PLoS One 8, e58936 (2013).

48. Coen, P., Xie, M., Clemens, J. & Murthy, M. Sensorimotor transformations underlying variability in song intensity during Drosophila courtship. Neuron 89, 629–644 (2016).

49. Sengupta, S., Chan, Y.-B., Palavicino-Maggio, C. B. & Kravitz, E. A. GABA transmission from mAL interneurons regulates aggression in Drosophila males. Proc. Natl. Acad. Sci. U. S. A. 119, e2117101119 (2022).

50. Sengupta, S. & Kravitz, E. A. Decoding sex differences: how GABA shapes Drosophila behavior. Curr. Opin. Insect Sci. 67, 101293 (2025).

51. Andrews, J. C., et al. Octopamine neuromodulation regulates Gr32a-linked aggression and courtship pathways in Drosophila males. PLoS Genet. 10, e1004356 (2014).

52. Certel, S. J., Savella, M. G., Schlegel, D. C. F. & Kravitz, E. A. Modulation of Drosophila male behavioral choice. Proc. Natl. Acad. Sci. U. S. A. 104, 4706–4711 (2007).

53. Sherer, L. M., et al. Octopamine neuron dependent aggression requires dVGLUT from dual-transmitting neurons. PLoS Genet. 16, e1008609 (2020).

54. Paulk, A. C., Kirszenblat, L., Zhou, Y. & van Swinderen, B. Closed-loop behavioral control increases coherence in the fly brain. J. Neurosci. 35, 10304–10315 (2015).

