## Supplementary figures and images for "High-Throughput Tracking of Freely Moving *Drosophila* Reveals Variations in Aggression and Courtship Behaviors"

### Supplementary Figure 1

**α- mCD8**

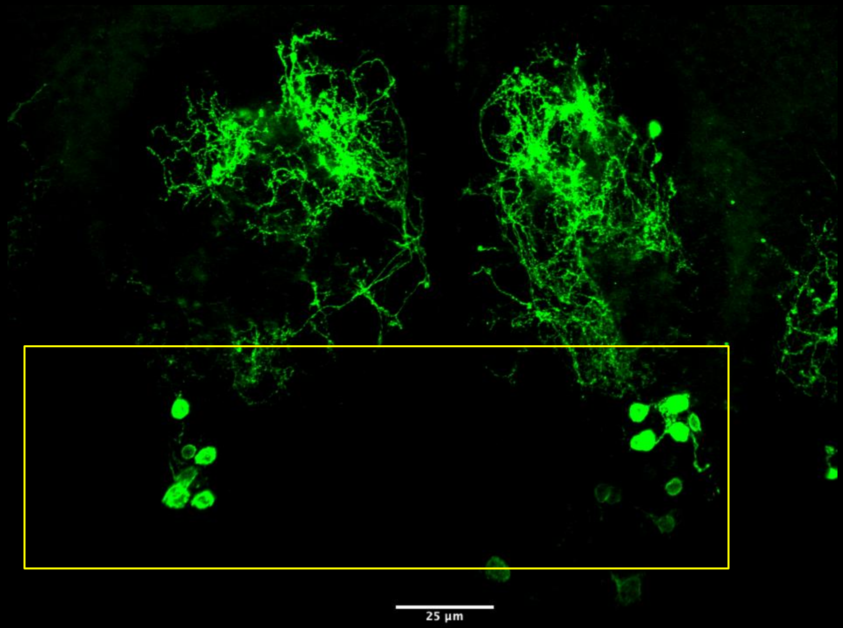


**α- Tdc2**

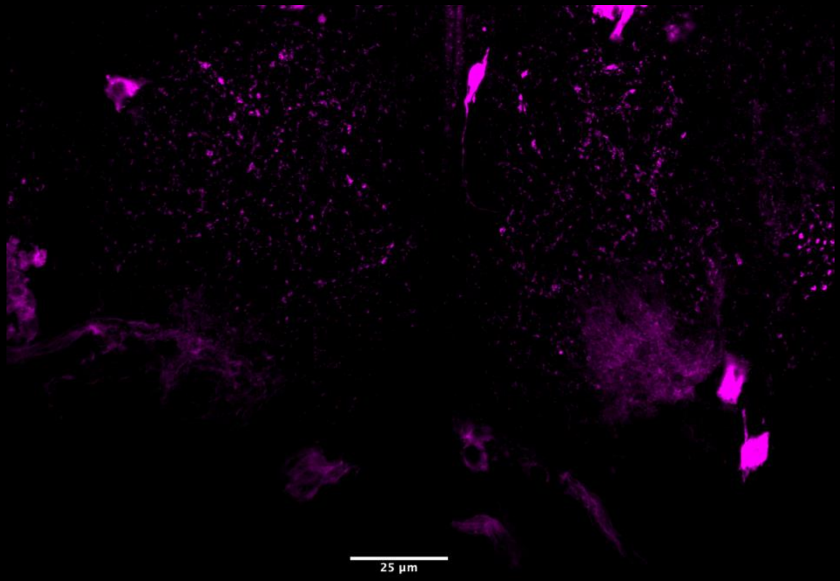


**Merge**

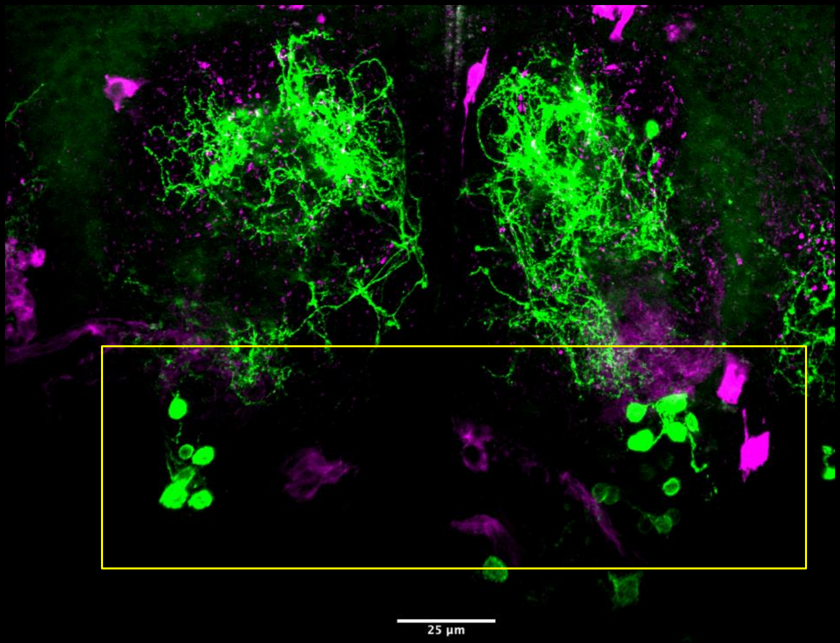
